# Functional analysis of chromatin-associated proteins in *Sordaria macrospora* reveals similar roles for RTT109 and ASF1 in development and DNA damage response

**DOI:** 10.1101/2023.10.17.562778

**Authors:** Jan Breuer, David Emanuel Antunes Ferreira, Mike Kramer, Jonas Bollermann, Minou Nowrousian

**Author notes:** author for correspondence: Minou Nowrousian Department of Molecular and Cellular Botany Ruhr University Bochum Universitätsstr. 150 44801 Bochum.

## Abstract

We performed a functional analysis of two potential partners of ASF1, a highly conserved histone chaperone that plays a crucial role in the sexual development and DNA damage resistance in the ascomycete *Sordaria macrospora*. ASF1 is known to be involved in nucleosome assembly and disassembly, binding histones H3 and H4 during transcription, replication and DNA repair and has direct and indirect roles in histone recycling and modification as well as DNA methylation, acting as a chromatin modifier hub for a large network of chromatin-associated proteins. Here, we functionally characterized two of these proteins, RTT109 and CHK2. RTT109 is a fungal-specific histone acetyltransferase, while CHK2 is an ortholog to PRD-4, a checkpoint kinase of *Neurospora crassa* that performs similar cell cycle checkpoint functions as yeast RAD53. Through the generation and characterization of deletion mutants, we discovered striking similarities between RTT109 and ASF1 in terms of their contributions to sexual development, histone acetylation and protection against DNA damage. Phenotypic observations revealed a developmental arrest at the same stage in Δrtt109 and Δasf1 strains, accompanied by a loss of H3K56 acetylation, as detected by western blot analysis. Deletion mutants of *rtt109* and *asf1* are sensitive to the DNA damaging agent MMS (methylmethane sulfonate), but not HU (hydroxyurea). In contrast, *chk2* mutants are fertile and resistant to MMS, but not HU. Our findings suggest a close functional association between ASF1 and RTT109 in the context of development, histone modification and DNA damage response, while indicating a role for CHK2 in separate pathways of the DNA damage response.

**Article summary:** In the filamentous fungus *Sordaria macrospora*, the conserved histone chaperone ASF1, which interacts with histones H3 and H4, was previously shown to be required for multicellular development and DNA damage response. Here, we have analyzed two additional chromatin-associated proteins. *rtt109* encodes a histone acetyltransferase, and deletion of the gene in *S. macrospora* results in a phenotype similar to that of a Δasf1 mutant, whereas *chk2* is involved in different aspects of the DNA damage response, but not in development.

## Introduction

The fruiting bodies of ascomycetes are some of the most intricate structures of the fungal kingdom. While fruiting body formation represents a significant developmental process in the fungal life cycle, our understanding of the specific factors that regulate this differentiation remains limited. *Sordaria macrospora*, a homothallic ascomycete, has proven to be an excellent model to gain insight into the genetic background of sexual development and multicellular development in general (1). *S. macrospora* offers multiple advantages as a model organism, such as a very fast life cycle as it generates its complex fruiting bodies, the perithecia, in under 7 days under laboratory conditions (2). Multiple approaches, from the investigation of mutants from random mutagenesis programs (3) to transcriptomics-based reverse genetics (4), have been used to study the complex genetic network of perithecia formation. Such research has yielded a lot of information about developmental genes, some of which are known to be conserved in higher eukaryotes and are in cases like the components of the STRIPAK complex (5) or the histone chaperone ASF1 (6, 7) even relevant for human diseases. ASF1 is known to interact with histones H3 and H4, both individually and as a dimer. It is involved in histone assembly and disassembly and was the first histone chaperone discovered to be involved in these processes during DNA replication, repair and transcription and was long thought to be the only one involved in all three processes (8). While ASF1 facilitates histone transfer and binds non-DNA bound H3-H4, it is not incorporated into the nucleosomes (9). During sexual development of *S. macrospora*, ASF1 has been shown to be essential for achieving fertility, and Δasf1 mutants arrest their life cycle at the protoperithecia stage (6). Effects on DNA methylation (10) and histone modification (11) have also been described in *S. macrospora asf1* deletion mutants. Since the structure of ASF1 is well characterized and no domains have been detected that would allow it to carry out such enzymatic reactions by itself (8), functions in DNA methylation and histone modification are likely to be carried out by interaction partners. Studies in *S. cerevisae* showed a loss of H3K56ac histone acetylation in *asf1* deletion mutants, and the enzyme identified as responsible for this modification was shown to be the HAT (histone acetyltransferase) Rtt109 (12). Indeed, an interaction between Asf1, Rtt109, Vps75 and the target histone H3 has been demonstrated in yeast, and it seems likely that acetylation occurs during the interaction of these proteins (13). Therefore, the loss of H3K56ac in *S. cerevisae* Δasf1 strains appears to be caused by disruption of the complex that allows Rtt109 to act on its target. The dependence of Rtt109 on interaction partners is one of its distinguishing features, as *in vitro* experiments with the *S. cerevisiae* proteins have shown that the HAT alone is unable to acetylate histones and only functions in the presence of at least one of its interaction partners Asf1 or Vps75, while the presence of both is required for full activity (12). Rtt109 is exclusively found in fungi (14) and is responsible for modifying specific sites on histone H3, namely K9, K27 and K56, as clearly demonstrated in yeast (15, 16). In addition to its role in histone acetylation, Rtt109 has been shown to be involved in a variety of processes in fungal model organisms. In *Fusarium graminearum*, it has been shown to be involved in perithecia morphogenesis, ascospore formation, conidiation and host plant infection (17). In *Aspergillus flavus* it is important for aflatoxin synthesis, virulence and growth (18), whereas *Aspergillus fumigatus* requires it for normal development and DNA damage response, as well as virulence (19). In *Neurospora crassa*, a close relative of *S. macrospora*, RTT109 has been shown to be necessary for the production of small RNAs (20). Therefore, RTT109 can be considered a major factor in many fungal models, suggesting its importance in fungi in general, while the fact that is only found in this group of organisms might make it an interesting target for anti-fungal drugs.

The histone acetylations generated by RTT109 are closely associated with newly synthesized histones and play a crucial role in the assembly of nucleosomes during DNA replication and repair processes (15). The involvement of H3K56ac in DNA damage repair, and thus the dependence of this process on Rtt109, has been demonstrated in *S. cerevisae rtt109* deletion mutants, which lack H3K56ac and are sensitive to the DNA double-strand break inducer methyl methanesulfonate (MMS) and the DNA replication inhibitor hydroxyurea (HU) (21). A similar relationship between ASF1, RTT109 and H3K56ac might exist in *S. macrospora*, since *asf1* deletion strains of this fungus show sensitivity to MMS and a reduction in H3K56ac (11). Therefore, we hypothesized that RTT109 might be the link between ASF1, histone acetylation and DNA damage repair in *S. macrospora* and chose SMAC_05078, the *S. macrospora* RTT109 homologue, as a target for functional characterization.

Another potential candidate, which might contribute to the reduced genomic stability of *asf1* mutants, could be the checkpoint kinase Rad53. Rad53 was shown to be important for DNA damage protection under the influence of MMS and HU in *S. cerevisiae* (22). *S. cerevisiae* Rad53 is known to physically and functionally interact with Asf1, and the activity of Rad53 is tightly regulated by its phosphorylation state. Hypophosphorylated Rad53 is known to be bound to Asf1 and maintained in an inactive state, while phosphorylation and release from the complex is a sign of active Rad53 (23). Deletion of *asf1* in budding yeast causes defects in DNA damage recovery and may be due to insufficient inactivation of Rad53 (24). In addition, *asf1* deletion mutants show similar sensitivities to MMS and HU as *rad53* deletion mutants (23). Rad53 is the functional equivalent of checkpoint kinases Chk2 in humans and PRD-4 in the filamentous fungus *Neurospora crassa*, since the corresponding genes can complement an *S. cerevisiae* Δrad53 mutant (25, 26). The *S. macrospora* ortholog to PRD-4/Chk2 is SMAC_00634 (called CHK2 hereafter), which was chosen for analysis to shed more light on the chromatin modifier network that contributes to the DNA damage protection functions of ASF1 in *S. macrospora*.

In this work, we created deletion mutants of *rtt109* and *chk2*, and analyzed their vegetative growth and potential for sexual development. We also performed genotoxic stress assays to check for similarities to *asf1* mutants. Furthermore, we determined the subcellular localization of the RTT109 protein by fluorescence microscopy and estimated the global amount of H3K56Ac in Δrtt109 by western blot analysis. We discuss our findings in the context of the roles of chromatin modifiers during DNA damage protection and regulation of complex multicellular development.

## Materials and Methods

### Strains, crosses and growth conditions

Strains utilized in this study can be found in Table S1 in File S1. These strains were cultivated at a temperature of 25 °C on either solid or liquid cornmeal medium (BMM) or complete medium (CM), following previously established protocols (27) (28). In order to facilitate genetic crosses, the spore color mutant fus was employed as a partner strain, enabling the identification of recombinant asci (3). Previously described methods were used to carry out the transformation of *S. macrospora* (28).

### Cloning procedures, oligonucleotides and plasmids

The oligonucleotides required for generating plasmids and conducting integration tests have been provided in Table S2 in File S1. Additionally, all the plasmids utilized in this study are listed in Table S3 in File S1. The plasmid containing the deletion sequence for *rtt109* was generated through yeast recombinant cloning (29). For assembling the deletion plasmid for *chk2,* and the complementation plasmids for *rtt109* and *chk2*, Golden Gate cloning methodology was employed (30, 31).

### Generation of *rtt109* and *SMAC_00634* deletion mutants and complementation strains

For conducting gene deletions, homologous recombination was utilized following established procedures (32). In brief, a deletion cassette was constructed, comprising a selection marker in the form of a hygromycin phosphotransferase gene under control of the constitutive P*gpd* promoter and T*trpC* terminator from *Aspergillus nidulans* (33), flanked by upstream and downstream regions of the target gene. This cassette was then cloned into a vector and transformed into the *S. macrospora* Δku70 strain after restriction and gel elution. Since KU70 is necessary for the ectopic integration of DNA, using a *ku70* defective strain facilitates homologous recombination (32). Transformants that exhibited resistance to hygromycin were selected and subsequent crosses were performed with the spore color mutant fus, since most *S. macrospora* transformants are heterokaryotic. Ascospores from such crosses were isolated to obtain homokaryotic deletion mutants and to enable the elimination of the Δku70 background. The resulting strains were verified through PCR. To complement the *rtt109* and *chk2* mutants, the assembled plasmid containing the respective gene fused to an eGFP tag, controlled by a P*gpd* promoter, was introduced into the corresponding strains via transformation. Ascospores were isolated from the transformants that reached fertility, allowing the acquisition of homokaryotic strains carrying the complementation construct.

### Growth tests and genotoxic stress assays

Growth tests were performed on BMM media and the growth front was documented every 24 hours. All tests were performed with three biological replicates. Statistical evaluation was conducted using a Student’s t-test. Phenotypic characterization involved observing and documenting the growth and fertility of the strains on BMM plates using a Stemi 2000-C stereomicroscope (Zeiss). To assess the resistance against methyl methanesulfonate (MMS), all strains were inoculated on BMM plates supplemented with 0.007% (v/v) MMS, and the growth was observed and documented over a period of 4 days. Similarly, resistance against hydroxyurea (HU) was evaluated on BMM plates containing 8 mM HU using the same approach.

### Microscopic analysis

To investigate the subcellular localization of RTT109-EGFP fusion proteins, the strains were cultivated on glass slides along with strains expressing H3-mRFP fusion proteins, as previously described (34). Heterokaryotic strains can therefore express both fluorescence-labelled proteins (35). Subsequently, light and fluorescence microscopy were performed using an AxioImager microscope (Zeiss), equipped with a Photometrix Cool SnapHQ camera (Roper Scientific). EGFP fluorescence was detected using a Chroma filter set 41017 (HQ470/40, HQ525/50, Q495lp), while mRFP fluorescence was detected using set 49008 (EG560/40x, ET630/75 m, T585lp). The acquired images were further processed using MetaMorph software (Molecular Devices).

### Western blot analysis

To evaluate differences in the global levels of H3K56ac between the Δrtt109 strains and the wild type, a western blot analysis was conducted using antibodies specific to this particular modification. The wild type strain SN1693, the Δrtt109 strain S60, and the corresponding complementation strain S60K1 were grown in liquid BMM medium (27) at 27 °C for 4 days. The mycelium was harvested by filtration, washed with PPP (28) and subsequently frozen in liquid nitrogen before being ground to a fine powder. Protein extraction was performed by mixing the powder with extraction buffer (50 mM Tris/HCl pH 7.5, 250 mM NaCl, 0.05 % NP-40, 0,3 % (v / v) Protease Inhibitor Cocktail Set IV (Calbiochem)). After 20 min centrifugation, the protein concentrations were determined using Bradford assays (36). Equal amounts of proteins were separated by SDS gel electrophoresis, and transferred onto a PVDF membrane through western blotting. H3K56ac-specific antibodies (Active Motif #39082) and H3-specific antibodies (Cell Signaling #9715) were used to detect the respective bands.

## Results

### Deletion of *rtt109* in *S. macrospora* leads to sterility and impairs vegetative growth

For the functional characterization of the histone acetyltransferase RTT109 in *S. macrospora*, a *ku70* deletion strain was transformed using a deletion cassette designed for the *rtt109* sequence. The deletion cassette consisted of a hygromycin phosphotransferase gene flanked by sequences homologous to the *rtt109* gene, enabling the replacement of the target gene with a resistance marker. Primary transformants that exhibited resistance to hygromycin were selected and subsequently crossed in order to obtain homokaryotic deletion mutants without the Δku70 background. The deletion of *rtt109* in the resulting strains was verified by PCR with primers designed to amplify a part of the gene of interest. The correct integration of the deletion cassette was verified by PCR (Figure S1 in File S1). Throughout the generation of the deletion strains, primary transformants did not exhibit any noticeable defects in sexual development and demonstrated the ability to achieve fertility. Since primary transformants are heterokaryotic in most cases, effects of gene deletions are often not directly obvious and sometimes appear only in homokaryotic strains. To obtain homokaryons, the primary transformants were crossed with the spore color mutant *fus* and the resulting homokaryotic ascospores were isolated. Homokaryotic Δrtt109 strains were found to be sterile (Table S4 in File S1). Microscopic examination confirmed a block in the sexual development of Δrtt109 at the stage of young protoperithecia (Figure 1). While the ascogonia and early protoperithecia were formed normally and at the expected time, no further structures of sexual development were observed. Even after a week, there was no developmental progression beyond that stage. The developmental effects of *rtt109* deletions in *S. macrospora* appeared to be even more severe than in other ascomycete model organisms such as *F. graminearum*, where a reduction in ascospore formation and aberrant perithecia have been previously documented (17). *rtt109* fused with a C-terminal eGFP tag was reintroduced into the validated deletion strains through ectopic integration to conduct complementation analysis and perform localization studies using fluorescence microscopy. Complementation strains regained fertility and were able to grow wild type-like sexual structures (Figure 1).

**Figure 1:**
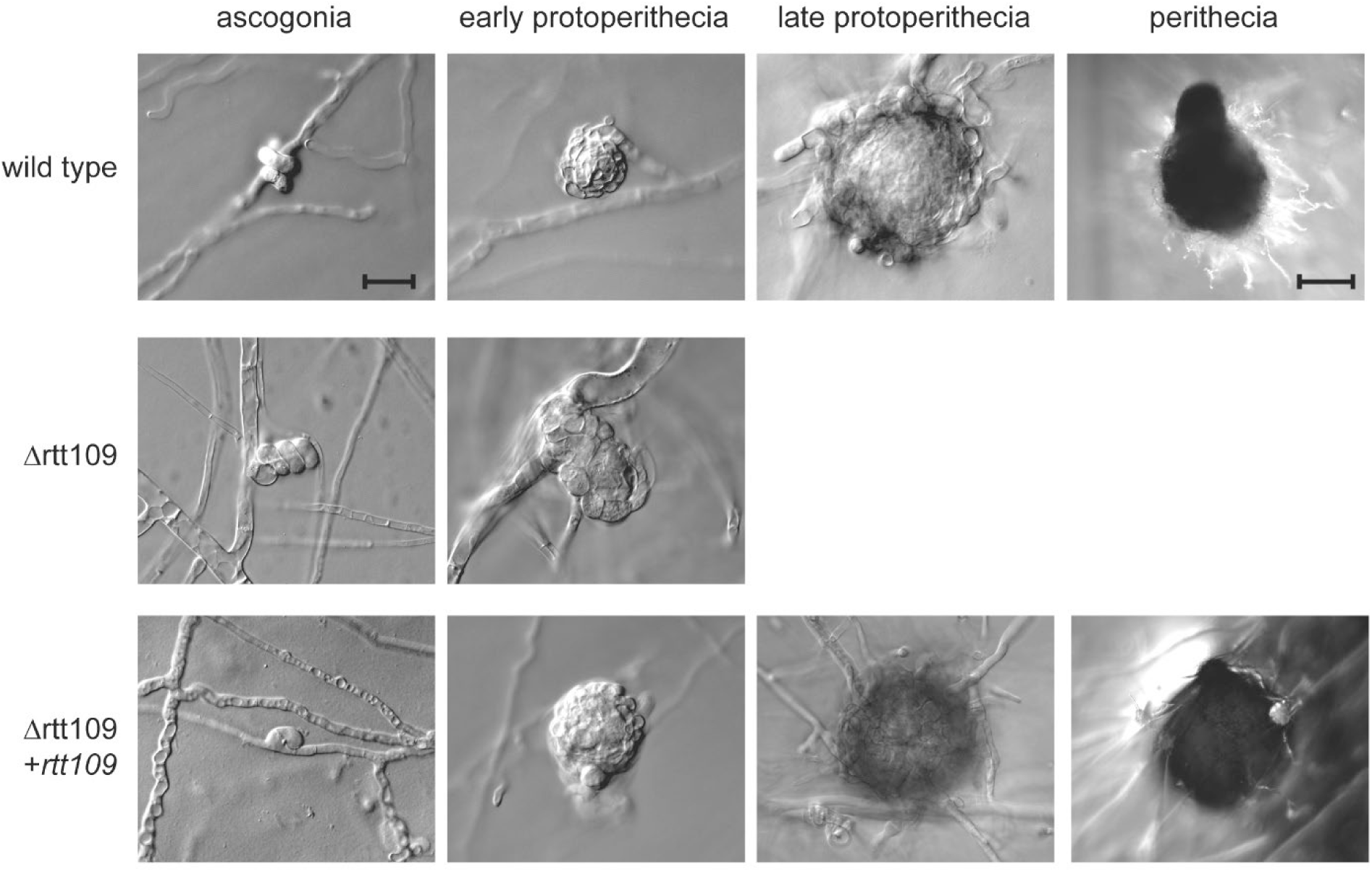
Development of *rtt109* deletion mutants compared to wild type and complementation strains. *S. macrospora* strains lacking the *rtt109* gene were found to exhibit impaired development with a block at the stage of early protoperithecia formation. Upon reintroduction of *rtt109*, normal life cycle progression was fully restored, as evidenced by the formation of late, melanized protoperithecia and the development of fully formed perithecia indistinguishable from those of the wild type strain. The scale bar for ascogonia and protoperithecia represents 20 µm, while the scale bar for perithecia represents 100 µm.

Deletion of *rtt109* also inhibited the vegetative growth rate of the respective mutants. Since RTT109 is suspected to be an interaction partner of ASF1, we documented the growth of *rtt109* deletion mutants, the respective complementation strain, and *asf1* deletion mutants to compare them to each other and to the wild type (Figure 2). Deletion of *rtt109* caused a visible reduction in vegetative growth rate, which was very similar to that of *asf1* deletion mutants, although the density of Δrtt109 mycelium appeared higher and more wild type-like. The growth rate of *rtt109* deletion strains was fully restored to wild type levels by reintroduction of the *rtt109* gene (Figure 2). We quantified this observation by measuring the progress of the growth front in 5 biological replicates and detected a reduction of vegetative growth speed of around 40 % in Δrtt109 and Δasf1 strains (Figure 3). The impairment of vegetative growth in *S. macrospora* Δrtt109 is consistent with results in *F. graminearum*, where corresponding deletion mutants were also reported to grow 40% slower than the wild type (17). This effect was completely reversed by complementation of the *rtt109* mutant and the respective strains exhibited a growth speed comparable to the wild type (Figure 3). In summary, the findings indicate that *rtt109* plays a crucial role in the sexual development of *S. macrospora* and is also significant for vegetative growth. The phenotype of *rtt109* deletion mutants appears highly similar to that of *asf1* deletion mutants, suggesting that indeed both function in the same pathway.

**Figure 2.**
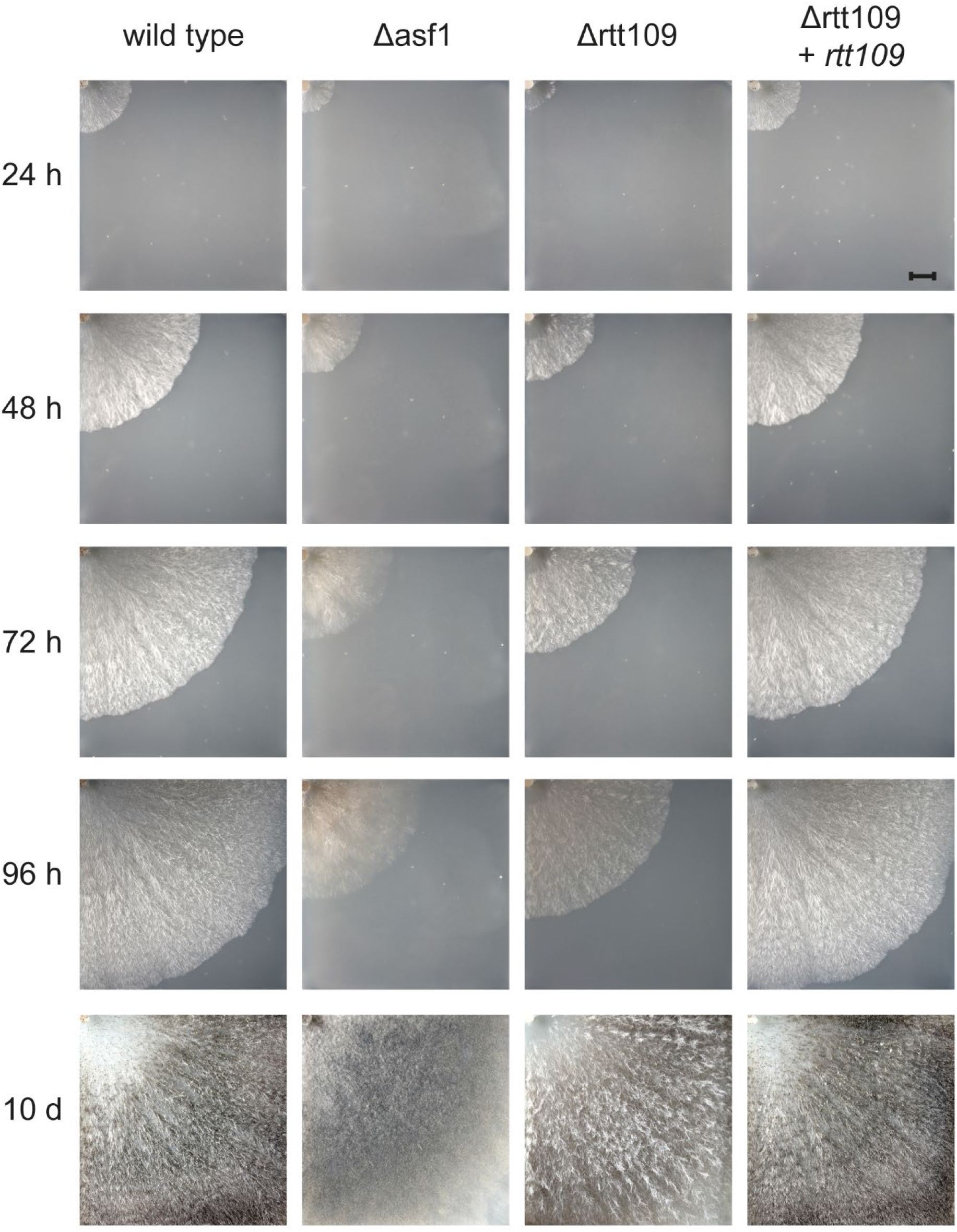
Comparison of vegetative growth of *rtt109* deletion and complementation strains with wild type and *asf1* deletion strains. Mycelial spread was observed over a period of 96 h and a final time after 10 d. While the wild type almost covered the entire plate in 96 h, *S. macrospora* Δrtt109 was significantly slower and grew as slowly as Δasf1 strains. This effect was reversed by reintroduction of *rtt109* in the *rtt109* deletion mutant, complementing the growth defect of the mutant. A visible difference between Δrtt109 and Δasf1 was the density of the mycelium. *S. macrospora* Δrtt109 appeared to grow as densely as the wild type, while Δasf1 appeared to be thinner overall. Scale bar represent 1 cm.

**Figure 3.**
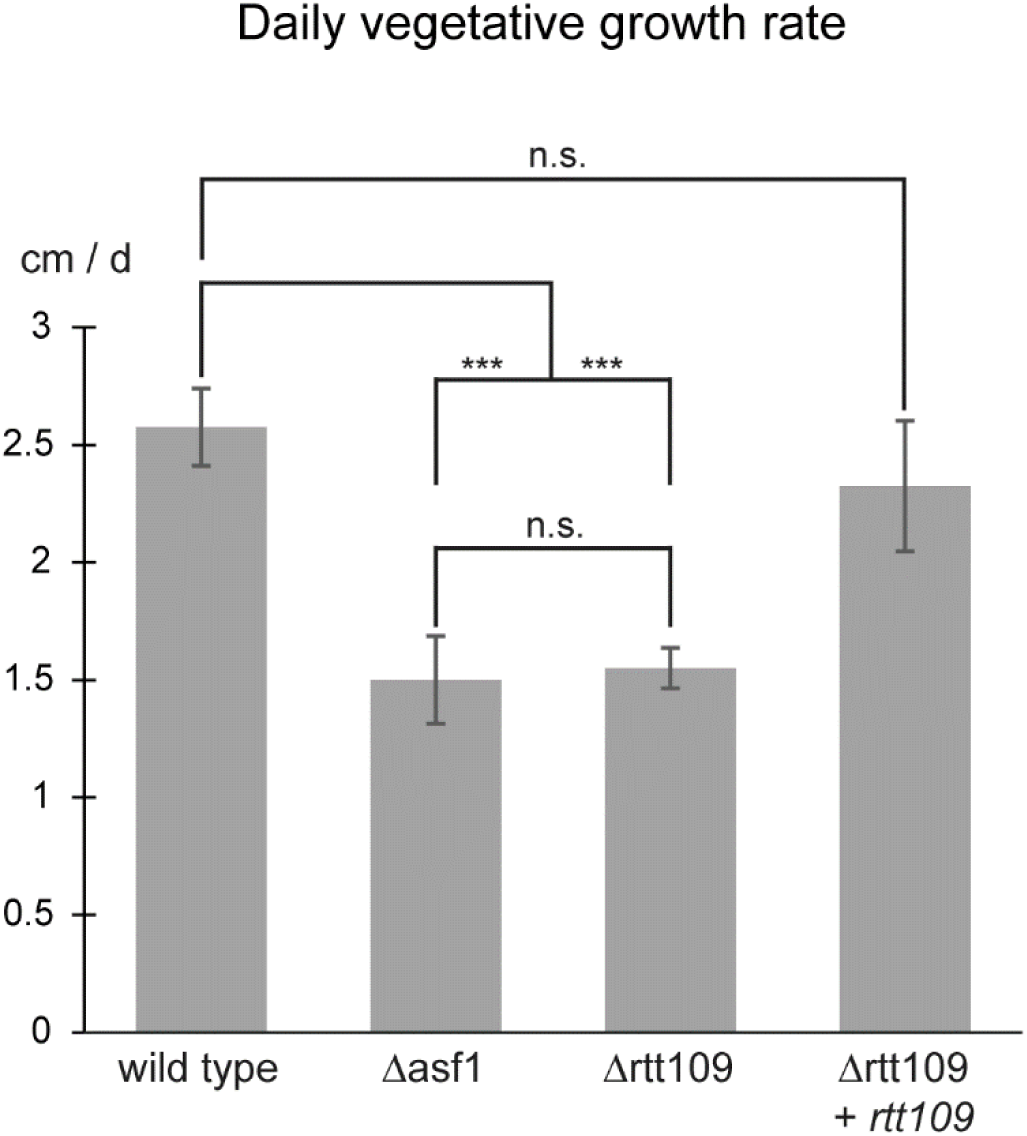
Quantification of the vegetative growth rates of *S. macrospora* Δrtt109, the respective complementation strain, Δasf1 and the wild type. While the wild type grew at about 2.5 cm per day, deletion of *rtt109* or *asf1* resulted in a significant decrease in growth rate to about 1.5 cm per day. Complementation of *rtt109* mutants restored the growth rate to wild type levels. Quantification was performed for 5 independent replicates and significance was assessed by Student’s t-test. *** = p-value < 0.001, n.s. = p-value > 0.05.

To further assess whether *asf1* and *rtt109* act in a similar manner during development, or whether there is a combinatorial effect in the absence of both genes, we attempted to cross Δasf1 and Δrtt109 strains to obtain double mutants. However, no fruiting bodies were formed in crosses of the two strains (Figure S2 in File S1). The inability of the mutants to cross with each other may be another indication that ASF1 and RTT109 act in a similar developmental pathway.

### *S. macrospora* Δrtt109 and Δasf1 react similarly to methyl methanesulfonate and hydroxyurea

In previous work, we discovered a sensitivity of *S. macrospora* Δasf1 strains to the DNA damaging agent MMS as well as a reduction in global H3K56ac levels in Δasf1 (11). The histone acetyltransferase RTT109 interacts with the chromatin modifier ASF1 in *Candida albicans* (13), *Schizosaccharomyces pombe* (37) and *S. cerevisiae* (14) and is involved in various mechanisms that ensure DNA stability during replication. Additionally, RTT109 is known as the primary enzyme responsible for H3K56 acetylation in *S. cerevisiae* (15). In the case of *A. fumigatus*, deletion of *rtt109* led to heightened sensitivity to DNA damaging agents such as MMS and HU (19). MMS induces DNA methyl adducts, which can cause double-strand breaks during replication (38), while HU inhibits the ribonucleotide reductase and leads to replication arrest by affecting dNTP supply (39). In our study, we exposed *S. macrospora* Δrtt109, the corresponding complementation strain, an *asf1* deletion mutant, and the wild type to MMS and HU. We found that the deletion of *rtt109* resulted in severe sensitivity to MMS, comparable to the sensitivity exhibited by the *asf1* mutant (Figure 4). While the wild type and complementation strains were able to grow relatively normally on BMM media containing MMS, the mutants failed to grow at all. Strikingly, we did not observe increased sensitivity against HU for the mutant strains. While aerial hyphae production appeared somewhat reduced, all strains were able to grow under HU stress (Figure 5). These observations contrast with the effects of *rtt109* deletions in *S. cerevisae* (15) and *A. fumigatus* (19), where *rtt109* mutants are sensitive to MMS as well as HU, suggesting the presence of additional factors in *S. macrospora* that respond to such specific stress conditions.

**Figure 4.**
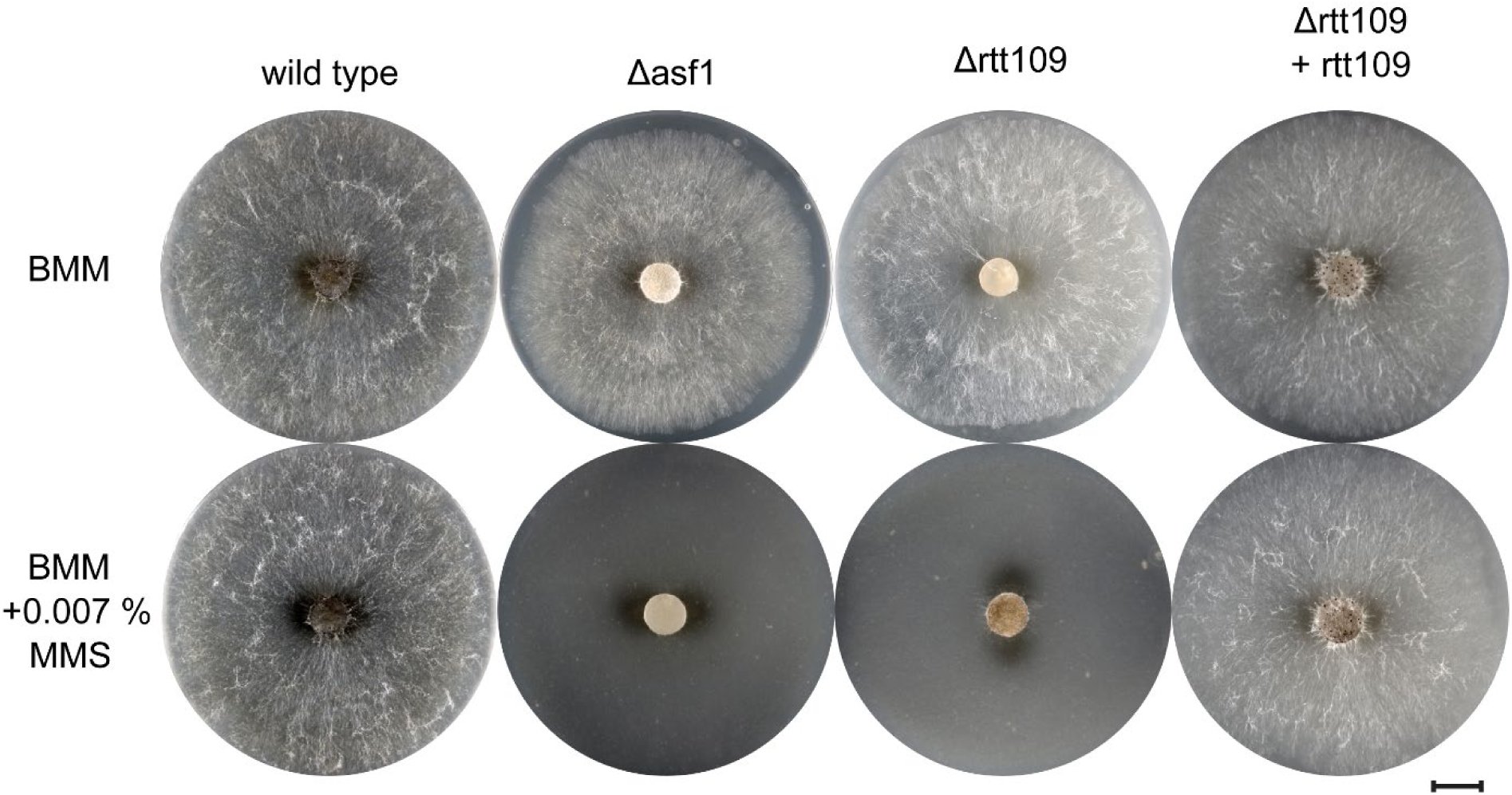
MMS sensitivity test for *S. macrospora* Δrtt109. The sensitivity of *S. macrospora rtt109* deletion mutants to the genotoxic compound MMS was observed after 4 days of growth on BMM media and BMM media supplemented with 0.007 % MMS. The addition of MMS completely halted the growth of the *rtt109* and *asf1* deletion mutants. When *rtt109* was reintroduced into Δrtt109, its ability to survive under MMS stress was restored. The scale bar provided represents 1 cm.

**Figure 5.**
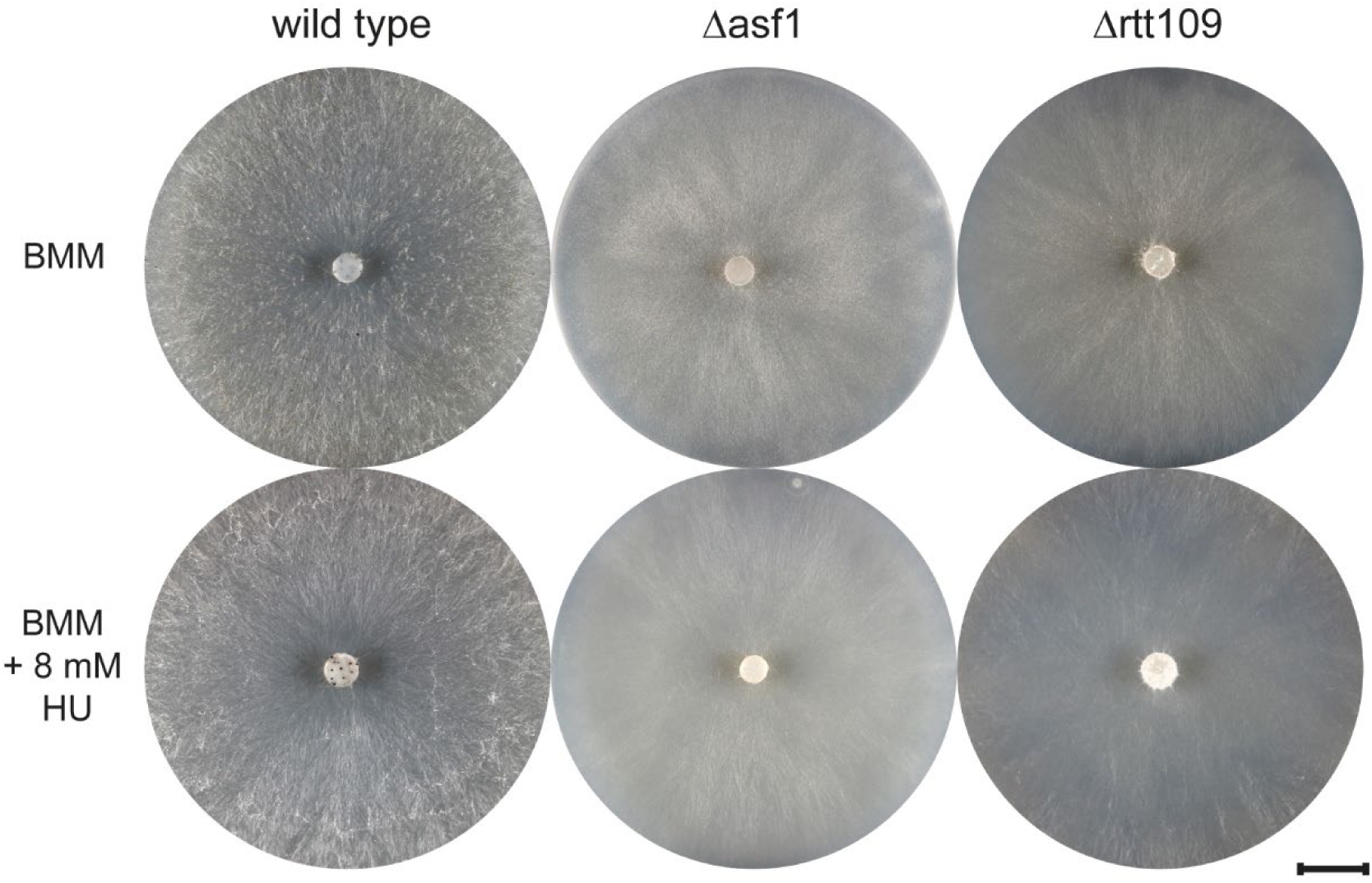
HU sensitivity test for *S. macrospora* Δrtt109. After a 4-day incubation on BMM media supplemented with 8 mM hydroxyurea, the sensitivity of *rtt109* deletion mutants was assessed. *S. macrospora* Δrtt109 exhibited no visible sensitivity to this DNA damaging agent, resembling the resistance observed in Δasf1. Scale bar represents 1 cm.

### *S. macrospora* RTT109 is localized in the nucleus

Given that RTT109 is described as a histone acetyltransferase, its targets are expected to be found within the nuclear compartment of the cell. Consequently, it is necessary for RTT109 to localize within the nucleus. Our complementation experiments with *rtt109* fused to an eGFP tag restored the phenotype of the deletion mutant, so a correct localization of the fusion protein was expected. To analyze the localization of RTT109, we employed fluorescence microscopy on the complementation strains. These strains were cultivated alongside a marker strain that expressed histone H3 fused to an mRFP tag. Since histones are primarily localized in the nucleus, this setup enabled us to perform co-localization analysis in the resulting heterokaryons (34). Through the detection of both green and red fluorescence in the same cellular compartment, we confirmed the co-localization of RTT109 and histone H3 (Figure 6). Based on these findings, we can conclude that RTT109 in *S. macrospora* is indeed localized within the nucleus.

**Figure 6.**
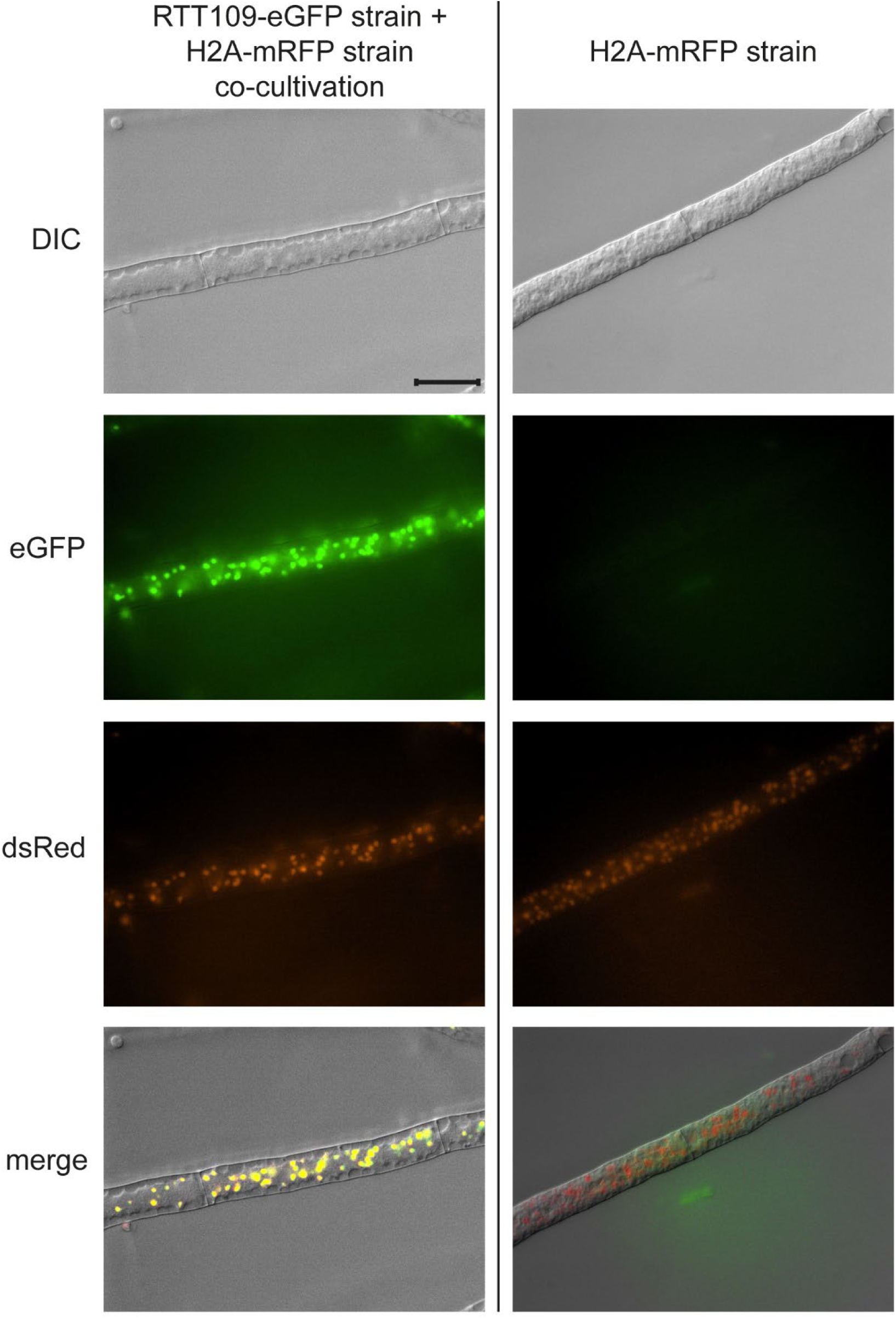
Localization analysis of RTT109 by fluorescence microscopy. RTT109 was expressed as a fusion protein with an eGFP tag and cultivated together with a strain expressing histone H3 fused to an mRFP tag. The visible colocalization of the green and red fluorescence indicates a nuclear localization of RTT109. No green autofluorescence was detectable in the reference strain expressing only H2A-mRFP. Scale bar represents 20 µm.

### The *rtt109* deletion leads to a loss of H3K56ac *S. macrospora*

RTT109 is widely recognized as the primary, and possibly exclusive, histone acetyltransferase responsible for the acetylation of H3K56 in fungal organisms (14, 15, 21). This specific histone modification has been demonstrated to play a crucial role in the cellular response to DNA damage induced by MMS (40). Therefore, we analyzed the levels of H3K56ac in the *S. macrospora* Δrtt109 mutant. To assess changes in the levels of global H3K56ac, we conducted western blot analysis in the *S. macrospora* Δrtt109 deletion mutant and compared the results to the wild type and the complemented mutant. Our findings revealed a loss of this histone modification in the deletion mutant (Figure 7, Figure S3 in File S1). The complementation strain exhibited a complete restoration of H3K56ac production, indicating the successful reinstatement of normal acetylation.

**Figure 7.**
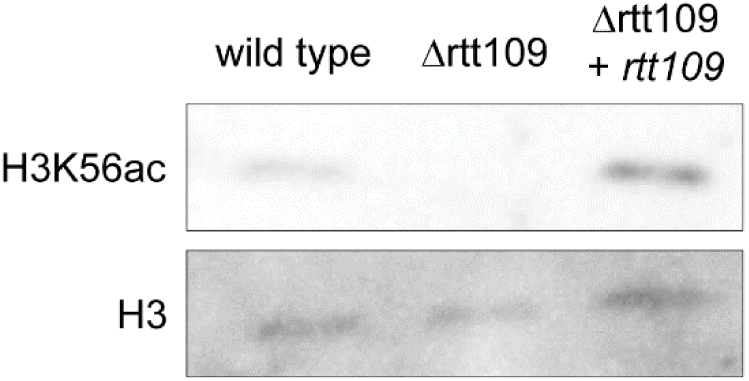
Assessment of H3K56ac levels in *S. macrospora* Δrtt109. The level of global H3K56ac was determined by comparing equal amounts of whole protein extracts from the wild type, Δrtt109, and the respective complementation strain by SDS-Page separation and western blotting with H3K56ac-specific antibodies. H3 antibodies were used to assess equal loading and comparable amounts of histone 3 in the protein extracts. In three biological replicates, no signal for H3K56ac was detectable in Δrtt109 strains. This effect was complemented by reintroduction of *rtt109* in the deletion mutant. Uncropped blots and the corresponding Coomassie gels are shown in Figure S3 in File S1.

### Deletion of *chk2*, a functional equivalent of *rad53*, has no effect on sexual development

The second potential partner for ASF1 during DNA damage protection we analyzed was the putative checkpoint kinase CHK2, which is orthologous to the checkpoint kinases PRD-4 from *N. crassa* and the human CHK2. It is the closest *S. macrospora* homolog to the *S. cerevisiae* checkpoint kinase Rad53 (Figure S4 in File S1). In contrast to the analyzed filamentous fungi and animals, there is a second homolog in *S. cerevisiae*, Dun1, which is more similar to PRD-4 and CHK2 proteins from animals both in sequence and domain structure than Rad53 (Figure S4B in File S1). However, PRD-4 from *N. crassa* and the human CHK2 were both shown to be able to complement an *S. cerevisiae* Δrad53 mutant and are therefore functionally equivalent to Rad53 (25, 26). Rad53 plays a critical role in maintaining genomic stability and regulating cell cycle arrest during DNA damage repair, as well as histone recycling in *S. cerevisiae* (41).

In yeast, the interaction between Asf1 and Rad53 is well documented, with Asf1 binding hypophosphorylated, inactive Rad53 and thus ensuring recovery from DNA damage repair (24). To investigate the role of CHK2 in *S. macrospora*, we deleted the *S. macrospora chk2* using homologous recombination. Hygromycin resistant primary transformants were crossed to obtain homokaryotic strains, and the deletion of *chk2* was confirmed by PCR (Figure S5 in File S1). Both primary transformants and homokaryotic deletion mutants showed normal growth and fruiting body development without any noticeable differences compared to the wild type. Vegetative growth appeared wild type-like (Figure 8) and Δchk2 strains achieved fertility within the expected timeframe. To further assess potential developmental defects, we examined the morphology of perithecia and asci in the mutant compared to the wild type and found no detectable differences (Figure 9). Thus, a deletion of *chk2* has no discernible effect on the development of *S. macrospora*.

**Figure 8:**
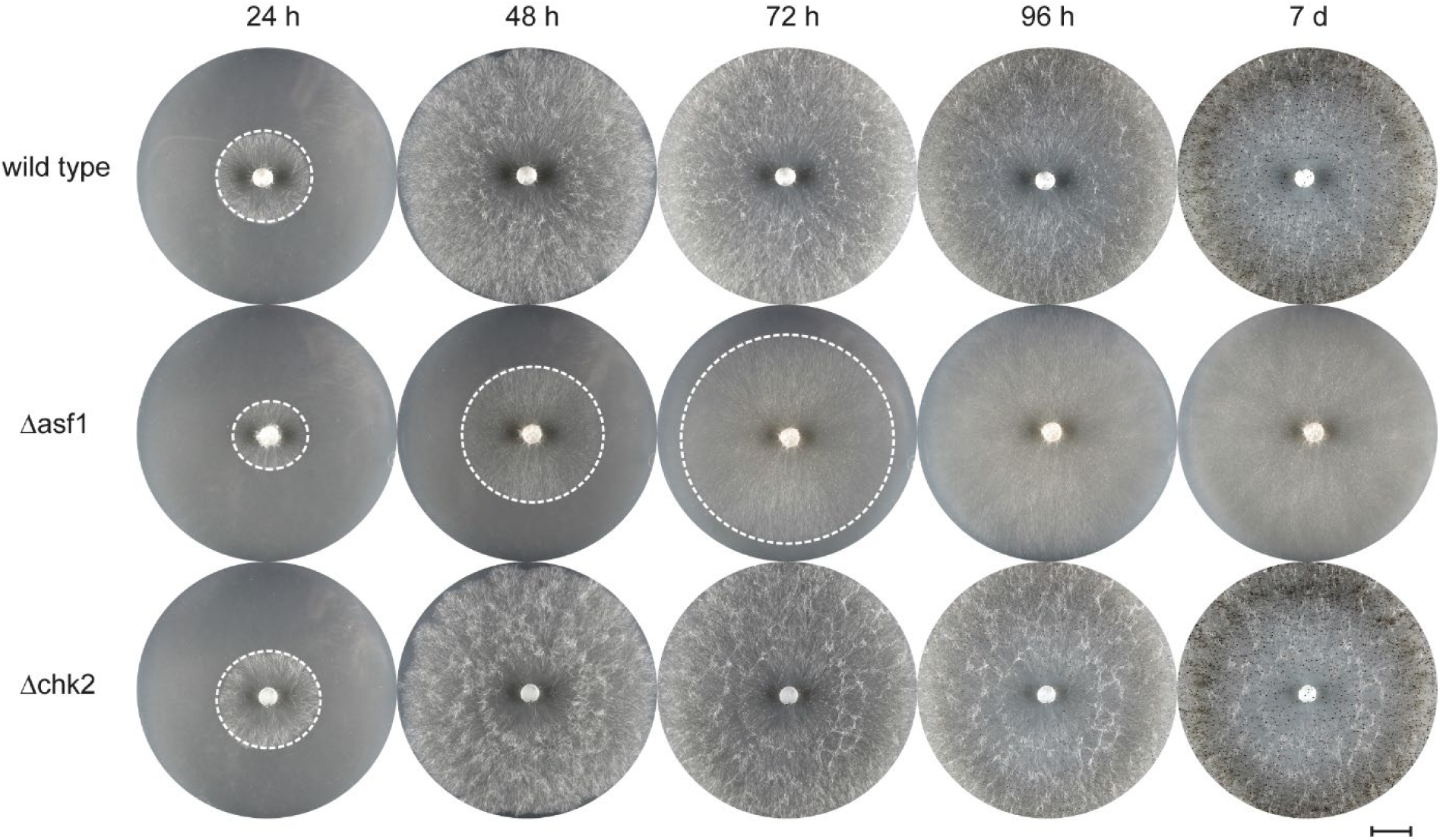
Growth comparison of *S. macrospora* Δchk2 with Δasf1 and the wild type. The growth and overall characteristics of the strains were monitored for 96 hours, with a final observation after 7 days on BMM media. There were no noticeable differences between the *chk2* deletion mutant and the wild type strain. Both strains exhibited similar growth rates and developed visible perithecia. The Δasf1 strain displayed significantly slower growth and failed to produce perithecia throughout the observation period. White dashed circles indicate the growth front. Scale bar represents 1 cm.

**Figure 9:**
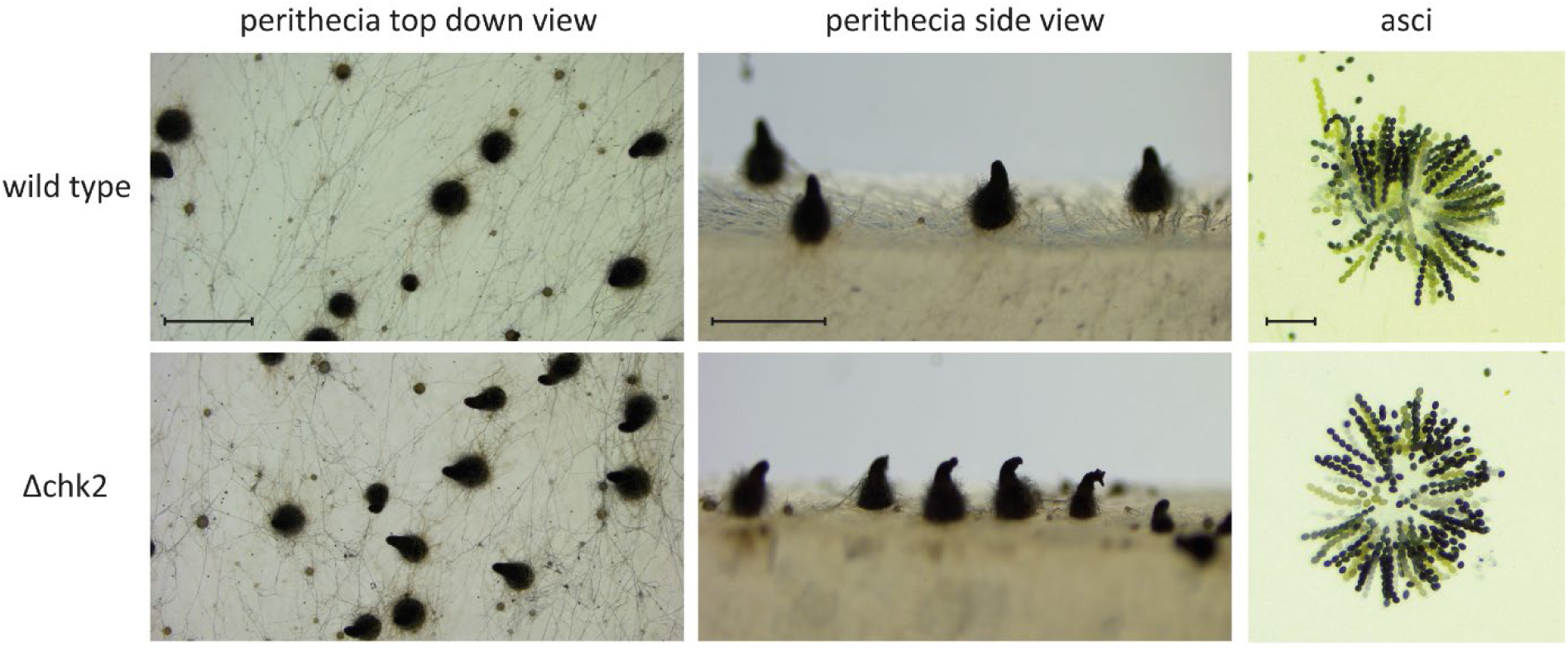
Morphological comparison of sexual structures generated by *S. macrospora* Δchk2 with those of the wild type. The *chk2* deletion mutant demonstrated the ability to undergo the complete life cycle of *S. macrospora* without any impairments. The fruiting bodies produced by the mutant exhibited no abnormalities in comparison to the wild type strain. The perithecia were formed and positioned in a normal manner and the overall morphology of the perithecia did not display any visible defects. Δchk2 strains generated wild type-like asci. Scale bars represent 1 mm for top down view, 500 µm for side view and 100 µm for asci.

### *S. macrospora* Δchk2 shows the opposite reaction to MMS and HU as Δrtt109 and Δasf1

Given the known involvement of RAD53 and its functional equivalent PRD-4 in DNA damage protection (42, 43), it was reasonable to expect a similar role for *S. macrospora* CHK2., Therefore, we conducted sensitivity tests with the genotoxic substances MMS and HU on the *S. macrospora chk2* deletion mutant. Surprisingly, the presence or absence of *chk2* did not have a noticeable impact on MMS resistance in *S. macrospora* (Figure 10), in contrast to findings with the *N. crassa* ortholog PRD-4 (25). This suggests that the relationship between ASF1 and CHK2 may not be significant in the context of MMS-induced stress in our model system. To evaluate the possibility that a CHK2 paralog might exist in *S. macrospora* that could substitute for CHK2 in the Δchk2 mutant and thus explain the lack of developmental phenotypes and the MMS resistance of the mutant, we performed BLASTP analysis (44) with CHK2. While several proteins showed partial similarity to CHK2, this similarity was restricted to the serine/threonine protein kinase domain, and none of the proteins contained the additional forkhead-associated domain that is present in CHK2 orthologs from *S. macrospora* and other fungi (Figures S4A, S6 and S7 in File S1). Thus, it appears unlikely that these putative kinases can substitute for CHK2 in the Δchk2 mutant.

**Figure 10:**
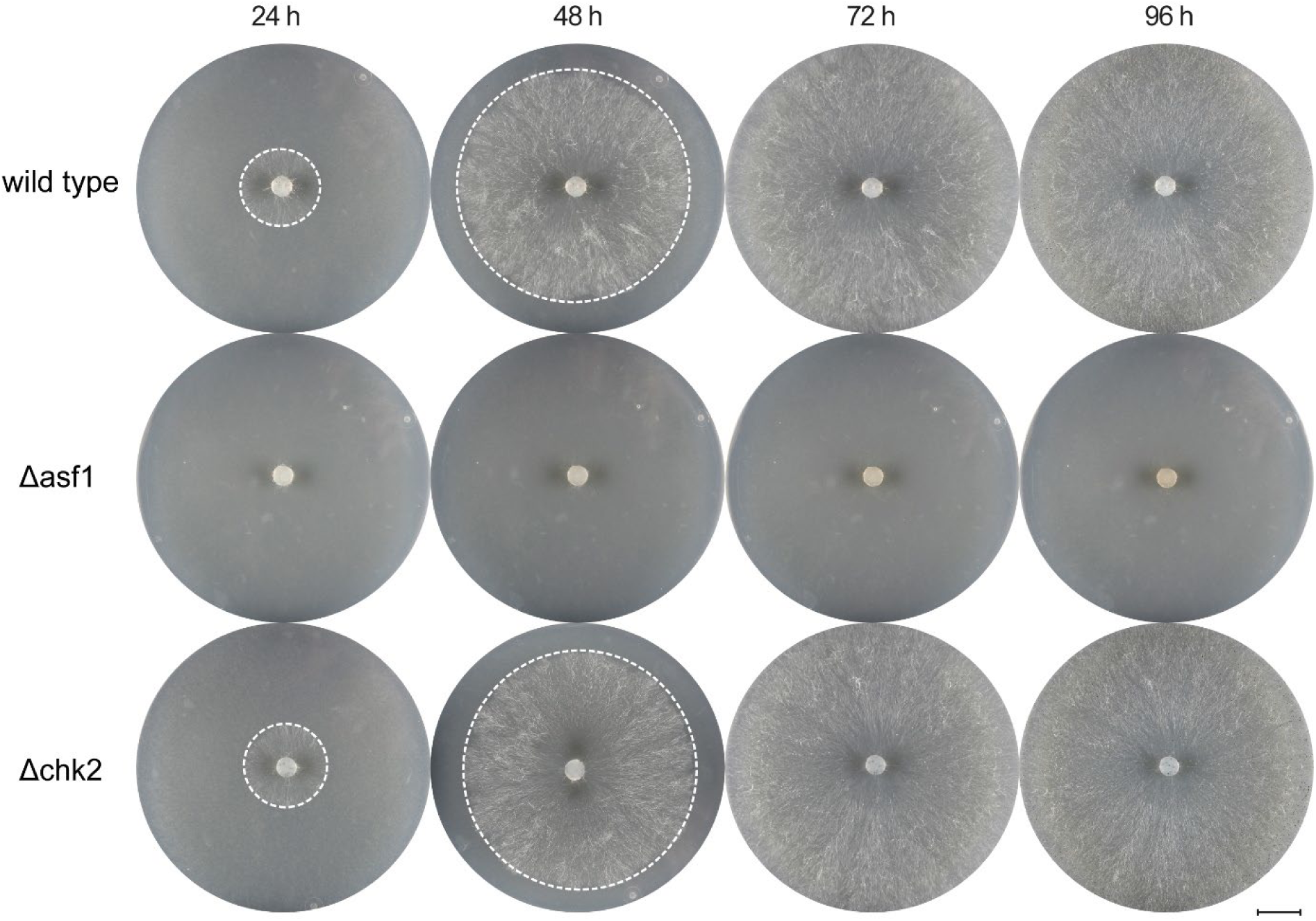
MMS sensitivity test for *S. macrospora* Δchk2. The growth of the strains was monitored on BMM media supplemented with 0.007% MMS for a duration of 96 hours. The *S. macrospora* Δchk2 strain did not display increased sensitivity to the DNA damaging agent compared to the wild type strain. Consistent with previous observations, the *asf1* deletion mutant exhibited a high level of sensitivity to MMS. White dashed circles indicate the growth front. Scale bar represents 1 cm.

In contrast to growth on MMS, under hydroxyurea stress, the deletion of *chk2* had a clear effect, unlike the deletion of *asf1* (Figure 11). Reintroduction of *chk2* into *S. macrospora* Δchk2 strains complemented the HU sensitivity phenotype (Table S5 in File S1). While all strains initially exhibited slow growth under HU stress without significant differences within the first 48 hours, Δchk2 mutants ceased growing after 48 h. In contrast, the wild type and *asf1* deletion strain proved to be resistant, consistent with previous observations comparing them to Δrtt109 strains. These results suggest that CHK2 plays a critical role in the DNA damage response pathway specific to coping with HU-induced stress, independent of ASF1 or RTT109.

**Figure 11:**
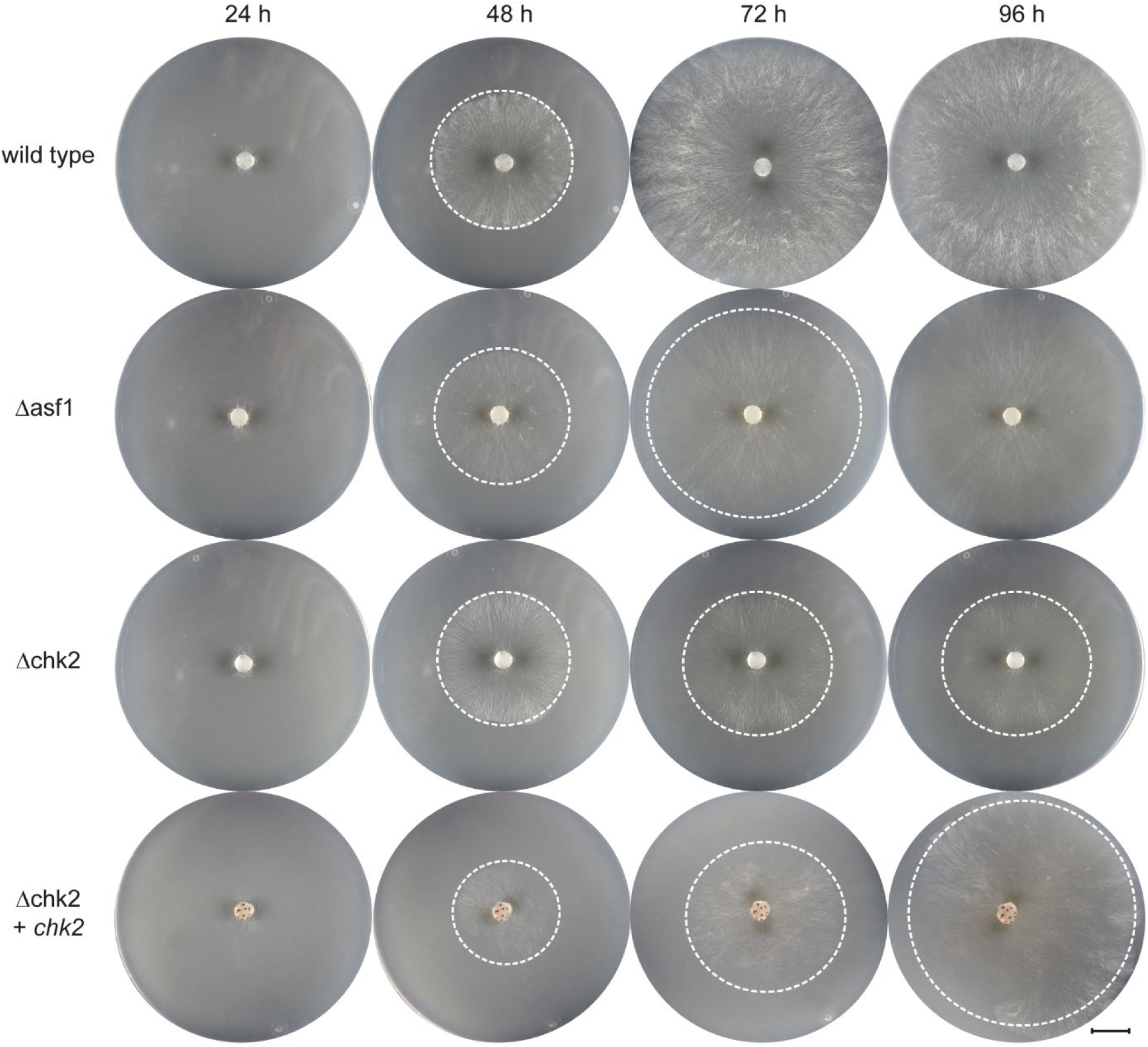
HU sensitivity test for *S. macrospora* Δchk2. The growth of the strains was observed on BMM media supplemented with 8 mM hydroxyurea for a period of 96 hours. In contrast to the observations with MMS, the *chk2* deletion mutant exhibited sensitivity to the DNA damaging agent hydroxyurea. While Δchk2 strains appeared to progress normally for the first 48 hours, they stopped growing after that time. Reintroduction of the *chk2* gene restored HU resistance. In contrast, the wild type and Δasf1 strains continued to grow under the same conditions. White dashed circles indicate the growth front. Scale bar represents 1 cm.

## Discussion

### RTT109 might be necessary for numerous functions of ASF1 during DNA damage protection and sexual development

In this study, we investigated the role of two potential partners of the histone chaperone ASF1 during DNA damage protection and sexual development in the ascomycete *S. macrospora*. Our findings suggest that the histone acetyltransferase RTT109 has significant functions under normal and DNA damage stress conditions. Deletion of *rtt109* resulted in strains that exhibited similarities to *asf1* deletions in terms of vegetative growth, sexual development, histone acetylation, and DNA damage response. These observations suggest a close relationship between ASF1 and RTT109 in these processes, as supported by their documented interactions in other fungal models like *C. albicans, S. pombe* and *S. cerevisae* (13, 14, 37). *S. macrospora rtt109*/*asf1* double mutants may be useful to determine if the two proteins act in the same pathway. A phenotype similar to single mutants with respect to growth and development should be observed in double mutants if RTT109 and ASF1 act in a cooperative manner during these processes. However, the single mutants proved to be unable to form sexual structures in genetic crosses. This could be the result of an effect on sexual development shared by both deletion strains, or the result of stress levels too high to allow proper crossing. The similarities between the respective deletion strains described raise the hypothesis that the severe phenotype observed in *S. macrospora* Δasf1 may be the consequence of the lack of interaction with RTT109. Studies in budding yeast have shown that Rtt109 relies on Asf1 to obtain histone H3 for acetylating K56 and the absence of Asf1 may nullify the functions of Rtt109 (45). In previous work, we demonstrated that sexual development in *S. macrospora* depends on the ability of ASF1 to bind histones and hypothesized that it acts as scaffold for H3-H4 interactions with other chromatin modifiers (11). The absence of such a scaffold might impair the ability of RTT109 to perform its functions in histone acetylation, which could cause the severe developmental defects in *asf1* deletion mutants. Thus, a reduction or improper positioning of H3K56ac may be the underlying issue in *S. macrospora* Δasf1 and proper acetylation of H3 may be crucial for sexual development. While overexpression experiments with *rtt109* in *S. macrospora* Δasf1 might provide information about the general level of H3K56ac necessary for development, positioning might still be problematic, and since ASF1 is thought to be the scaffold necessary for RTT109 to function at its full extent (12), higher RTT109 levels might not even correlate with higher H3K56ac levels in *asf1* deletion mutants. However, given the wide range of putative and proven functions of ASF1 (46), it may be challenging to pinpoint its role during *S. macrospora* development to a single interaction partner and its effect on H3K56ac. Interestingly, the DNA damage response functions of ASF1 do not seem to depend on its interaction with histones in the same way as sexual development, although ASF1 variants unable to bind histones showed a similar reduction of H3K56ac as full deletion mutants (11). However, our results on the sensitivity of *rtt109* deletion mutants to MMS show a dependence on RTT109 for DNA damage protection and therefore probably for the establishment of H3K56ac. More sensitive assays such as ChIP-seq are needed to fully quantify the potential differences in H3K56ac levels between *S. macrospora* Δasf1 and Δrtt109, and our results indicate that deletion of the HAT causes an even greater decrease, or even complete loss in H3K56 acetylation levels than the loss of a putative scaffold. In *S. cerevisiae*, Rtt109 is known to interact not only with Asf1 but also with another histone chaperone, Vps75, which also binds H3 and H4 (14). Although a fully functional Asf1-Vps75-Rtt109 complex is known to be necessary for establishing correct acetylation patterns in budding yeast (47), the mere presence of ASF1, even without histone binding ability, might be sufficient to enable lower-level activity of RTT109, providing some form of DNA damage protection. Therefore, the severe developmental defects observed in *S. macrospora* Δasf1, persistent when expressing ASF1 variants unable to bind histones, may have roots not only in the misregulation of developmental processes, but also in the accumulation of underlying DNA damage events, compromising the respective strains as a whole. Such DNA damage accumulation might be too insignificant to turn up during a sensitivity assay, but could potentially lead to disturbances during tightly regulated processes, such as sexual development and fruiting body formation. While this hypothesis requires further investigation, such as interaction studies between RTT109, ASF1 and VPS75, the role of RTT109 during sexual development and the DNA damage response in *S. macrospora* appears to be quite fundamental. As the primary facilitator of H3K56ac in fungi (15), RTT109 is clearly essential for survival under MMS conditions, which can induce double-strand breaks during replication (38). Furthermore, RTT109 is crucial during the formation of complex sexual structures, either by providing the necessary genomic stability for such processes or by ensuring the correct transcription of important genes through the establishment of proper acetylation patterns. Surprisingly, the deletion of *rtt109* did not increase the sensitivity of *S. macrospora* to hydroxyurea, despite its well-known DNA damaging properties that cause replication fork arrest by disrupting dNTP availability (39). In contrast, other fungal model organisms like *A. fumigatus* and *S. cerevisiae* have been shown to exhibit heightened sensitivity to HU when *rtt109* is deleted (19, 48).

### CHK2 is not essential for sexual development, but provides resistance to genotoxic stress independent of ASF1

The second putative partner of ASF1 during DNA damage protection we investigated was CHK2. CHK2 is the ortholog of *N. crassa* PRD-4, which can complement a Δrad53 mutant with respect to its function in DNA damage response in *S. cerevisiae* (25). Deletion of *chk2* did not result in any noticeable defects in vegetative growth or developmental processes, suggesting its negligible role in the formation of fruiting bodies. Furthermore, the mutants lacking *chk2* did not exhibit increased sensitivity to MMS-induced DNA damage stress, unlike the highly sensitive *asf1* and *rtt109* mutants. This indicates that CHK2 operates in a distinct damage response pathway from ASF1 and RTT109 under these specific conditions. However, we discovered a role for CHK2 in response to a different type of genotoxic stress, as the mutants showed sensitivity to the replication fork stressor hydroxyurea, while the *asf1* and *rtt109* mutants did not display such sensitivity. The budding yeast equivalent, Rad53, is known to be important for restarting inhibited replication forks (49), so it is reasonable to assume that the replication fork stalling caused by hydroxyurea cannot be efficiently rescued in the absence of Rad53, or in this case CHK2 in *S. macrospora*. Since Δasf1 strains are not inhibited under hydroxyurea stress, this suggests that CHK2 is involved in a genomic protection system that functions independently of ASF1. Notably, this observation appears to be specific for *S. macrospora*, as an *S. cerevisiae* Δasf1 mutant is sensitive to hydroxyurea (50). The precise functions of CHK2 and its relationship with ASF1 in its broader chromatin modification network are more challenging to elucidate, given the specific roles fulfilled by homologous checkpoint kinases like PRD-4 or functional equivalents like RAD53 in their respective native organisms. However, it can be inferred that any connection between ASF1 and CHK2 in DNA damage protection and sexual development is of minor importance. Additionally, the diverse functions of ASF1 in *S. macrospora* do not seem to include an active role in the response to hydroxyurea-induced stress.

In conclusion, our study revealed an essential role of the histone acetyltransferase RTT109 during sexual development, vegetative growth, histone modification and genotoxic stress response in the ascomycete *S. macrospora*. The phenotypic aberrations in *rtt109* deletion strains closely resemble those previously observed in *asf1* deletion mutants, indicating a strong correlation between the functions of ASF1 and RTT109. CHK2, another potential component of the ASF1-mediated chromatin modifier network, appears to be unrelated to these processes but is instead involved in a distinct DNA damage response system that operates independently of ASF1.

## Data Availability Statement

Strains and plasmids are available upon request. The authors affirm that all data necessary for confirming the conclusions of the article are present within the article, figures, and tables.

## Acknowledgements

The authors would like to thank Silke Nimtz for excellent technical assistance and Christopher Grefen for support at the Department of Molecular and Cellular Botany. This work was funded by the German Research Foundation (DFG, grant NO407/7-2 to MN).

## Notes

### Competing Interest Statement

The authors have declared no competing interest.

### Summary of Updates

Figures 6 and 7 revised, text sections on double mutants as well as homologs of chk2 added.

