## Supplemental File S1 for "Functional analysis of chromatin-associated proteins in *Sordaria macrospora* reveals similar roles for RTT109 and ASF1 in development and DNA damage response"

**File S1: Supplemental data for manuscript**

**"Functional analysis of chromatin-associated proteins in *Sordaria macrospora* reveals similar roles for RTT109 and ASF1 in development and DNA damage response"**

This document contains the following supplemental data:

Tables S1-S5

Figures S1-S7

**Table S1.** *S. macrospora* strains used in this study.

| **Name of strain** | **Genotype** | **Comments** |
|---|---|---|
| SN1693 | Wild type | Reference strain |
| SN1891 | Fus1-1 | Spore color mutant |
| SN1983 | Δasf1::*hph* | *asf1* deletion mutant, sterile |
| SN1835 | Δku70*::nat* | *ku70* mutant, defect in NHEJ, fertile |
| S60 | Δrtt109*::hph* | *rtt109* deletion mutant, sterile |
| S67 | Δrtt109*::hph* | *rtt109* deletion mutant, sterile |
| S69 | Δrtt109*::hph* | *rtt109* deletion mutant, sterile |
| S166 | Δrtt109*::hph* | *rtt109* deletion mutant, sterile |
| S60K1 | Δrtt109*::hph* + pCompRTT109-C-eGFP | *rtt109* deletion mutant with integrated complementation plasmid, fertile |
| S67K1 | Δrtt109*::hph* + pCompRTT109-C-eGFP | *rtt109* deletion mutant with integrated complementation plasmid, fertile |
| S69K1 | Δrtt109*::hph* + pCompRTT109-C-eGFP | *rtt109* deletion mutant with integrated complementation plasmid, fertile |
| DBT2-1-4-4 | Δchk2*::hph* | *SMAC_00634* deletion mutant, fertile |
| DBT2-1-4-1 | Δchk2*::hph* | *SMAC_00634* deletion mutant, fertile |
| RL2772 | Wild type + *H2A-mRFP::nat* | Nuclear marker strain, mRFP |
| SN7182 | Δchk2*::hph* + pSMAC_00634_egfp | *chk2* deletion mutant with integrated complementation plasmid, fertile |
| SN7186 | Δchk2*::hph* + pSMAC_00634_egfp | *chk2* deletion mutant with integrated complementation plasmid, fertile |
| SN7190 | Δchk2*::hph* + pSMAC_00634_egfp | *chk2* deletion mutant with integrated complementation plasmid, fertile |
| SN7195 | Δchk2*::hph* + pSMAC_00634_egfp | *chk2* deletion mutant with integrated complementation plasmid, fertile |
| SN7198 | Δchk2*::hph* + pSMAC_00634_egfp | *chk2* deletion mutant with integrated complementation plasmid, fertile |
| SN7199 | Δchk2*::hph* + pSMAC_00634_egfp | *chk2* deletion mutant with integrated complementation plasmid, fertile |
| SN7201 | Δchk2*::hph* + pSMAC_00634_egfp | *chk2* deletion mutant with integrated complementation plasmid, fertile |
| SN7202 | Δchk2*::hph* + pSMAC_00634_egfp | *chk2* deletion mutant with integrated complementation plasmid, fertile |
| SN7211 | Δchk2*::hph* + pSMAC_00634_egfp | *chk2* deletion mutant with integrated complementation plasmid, fertile |
| SN7215 | Δchk2*::hph* + pSMAC_00634_egfp | *chk2* deletion mutant with integrated complementation plasmid, fertile |
| SN7217 | Δchk2*::hph* + pSMAC_00634_egfp | *chk2* deletion mutant with integrated complementation plasmid, fertile |
| SN7220 | Δchk2*::hph* + pSMAC_00634_egfp | *chk2* deletion mutant with integrated complementation plasmid, fertile |

**Table S2.** Oligonucleotides used in this study.

| Primer | Sequence 5'-3' | remarks |
|---|---|---|
| SMAC_05078_ko1 | GCGCGCGTAATACGACTCACTATAGGGAATTC GCTAGCCGATTGAAGTTAGCTCAA | Amplification of *rtt109* flanks to construct pKO_SMAC_05078 |
| SMAC_05078_ko2 | GGGCAAAGGAATAGGGTTCCGTTCTTAGCAGT ATGTCTTGTCGGGTTTCT | Amplification of *rtt109* flanks to construct pKO_SMAC_05078 |
| SMAC_05078_ko3 | AAAAATGCTCCTTCAATATCAGTTTAAGATCA GCGAGGCCGGCATGGG | Amplification of *rtt109* flanks to construct pKO_SMAC_05078 |
| SMAC_05078_ko4 | TACGCCAAGCGCGCAATTAACCCTCAGAATTC ACAAGAGGATATGATGGGAAGGAACATACGC | Amplification of *rtt109* flanks to construct pKO_SMAC_05078 |
| GG_rtt109_comp_fw | GGCTACGGTCTCATGGTATGGCATCCTCGTCG TCG | Amplification of *rtt109* to construct pCompRTT109-C-eGFP |
| GG_rtt109_comp_rev | GGCTACGGTCTCGTACGTTAAGCTGATGTCTTG GGCTTC | Amplification of *rtt109* to construct pCompRTT109-C-eGFP |
| SMAC_05078_IP_fw | AGTCATTGCCTTCGCCATTGAG | Verification of *rtt109* deletion |
| SMAC_05078_IP_rv | GGTCAGCTGTGAAAGAGGCC | Verification of *rtt109* deletion |
| GG_3flank_fw | GGCTACGGTCTCCAGTTCATTAACACTTGCC | Amplification of *SMAC_00634* flanks to construct pKO_00634GG |
| GG_3flank_rv | GGCTACGGTCTCTTACGAGCCTATATCCAACTAACC | Amplification of *SMAC_00634* flanks to construct pKO_00634GG |
| GG_5flank_fw | GGCTACGGTCTCGTGGTGGATCCGACAATCCCTGGA GGACTATC | Amplification of *SMAC_00634* flanks to construct pKO_00634GG |
| GG_5flank_rv | GGCTACGGTCTCAATTGGCTGCTGGGATTGTAGG | Amplification of *SMAC_00634* flanks to construct pKO_00634GG |
| GG_hph_fw | GGCTACGGTCTCACAATTCGTGATAGGGCCCTC | Amplification of *hph* sequence to construct pKO_00634GG |
| GG_hph_rv | GGCTACGGTCTCCAACTCCCTAGCAACTGATATTGAAG GAGC | Amplification of *hph* sequence to construct pKO_00634GG |
| SMAC_00634_Int_fw | GCACGAGTTGGACAAGACCAAG | Verification of *SMAC_00634* deletion |
| SMAC_00634_Int_rv | GATCCGAAAGAGAATAAGGGAAGTCG | Verification of *SMAC_00634* deletion |
| hph1MN | CGATGGCTGTGTAGAAGTACTCGC | Verification of correct *hph* sequence insertion |
| hph2MN | ATCCGCCTGGACGACTAAACCAA | Verification of correct *hph* sequence insertion |
| KO_rtt109_left_fw | CCGTTTCCACTCTCGAATCGC | Verification of correct 5' *rtt109* deletion cassette insertion |
| KO_rtt109_right_rev | GTACCTCAAGACGGTCAAGGG | Verification of correct 3' *rtt109* deletion cassette insertion |
| SMAC_00634_L_1 | CGTGCTGATGGTCAGGGTTAT | Verification of correct 5' *chk2* deletion cassette insertion |
| SMAC_00634_L_5 | CATGATGCTCATGAAAGACAGCAC | Verification of correct 3' *chk2* deletion cassette insertion |

**Table S3.** Plasmids used in this study.

| Plasmid | Characteristics | Comments |
|---|---|---|
| pRS426 | 5715 bp, f1_ori, pBR322_ori, 2 micron_ori, P*ura3::ura3*, P*amp::bla*, *lacZ* | Backbone for cloning of pKO_SMAC_05078 |
| pGG-N_3xFLAG | 5,964 bp, pUC_ori, P*trpC::Nat*, P*gpd*, 3xFlag, *LacZa*, T*rpC*, P*amp::bla* | Backbone for cloning of pKO_SMAC_00634 |
| pGG-C-eGFP | 6,6 kb, P*trpC::nat1*, P*gpd*, *LacZa*, *eGFP*, T*trpC*, *bla*, pUC ori | Backbone for cloning of pCompRTT109-C-eGFP |
| pKO_RTT109 | 8,2 kb, pRS426 derivative, 3' and 5' flanks of *SMAC_05078*, *hph* | Deletion construct for *rtt109* |
| pKO_00634GG | 8,8 kb, pGG-N_3xFLAG derivative, 3' and 5' flanks of *SMAC_00634*, *hph* | Deletion construct for *SMAC_00634* |
| pCompRTT109-C-eGFP | 7,9 kb, pGG-C-eGFP derivative, *SMAC_05078-eGFP*, *nat* | Complementation construct for Δrtt109 strains with eGFP tag |
| pSMAC_00634_egfp | 11,2 kb pGG-C-eGFP derivative, *SMAC_000634-eGFP*, *nat* | Complementation construct for Δchk2 strains with eGFP tag |

**Table S4.** Fertility rates for primary transformants and homokaryotic spore isolates of *rtt109* deletion mutants and complementation strains.

| Genotype | Primary transformants | Fertile transformants | % fertile | Spore isolates | Fertile isolates | % fertile isolates |
|---|---|---|---|---|---|---|
| Δrtt109 | 18 | 18 | 100 % | 23 | 0 | 0 % |
| Δrtt109 + *rtt109* | 9 | 9 | 100 % | 44 | 32 | 72 % |

**Table S5.** HU sensitivity of *S. macrospora* Δchk2 and *chk2* complementation strains

| Strain | Genotype | Reaction to Hydroxyurea |
|---|---|---|
| DBT2-1-4-4 | Δchk2 | sensitive |
| DBT2-1-4-1 | Δchk2 | sensitive |
| SN7182 | Δchk2 + *chk2* | resistant |
| SN7186 | Δchk2 + *chk2* | resistant |
| SN7190 | Δchk2 + *chk2* | resistant |
| SN7195 | Δchk2 + *chk2* | resistant |
| SN7198 | Δchk2 + *chk2* | resistant |
| SN7199 | Δchk2 + *chk2* | resistant |
| SN7201 | Δchk2 + *chk2* | resistant |
| SN7202 | Δchk2 + *chk2* | resistant |
| SN7211 | Δchk2 + *chk2* | resistant |
| SN7215 | Δchk2 + *chk2* | resistant |
| SN7217 | Δchk2 + *chk2* | resistant |
| SN7220 | Δchk2 + *chk2* | resistant |

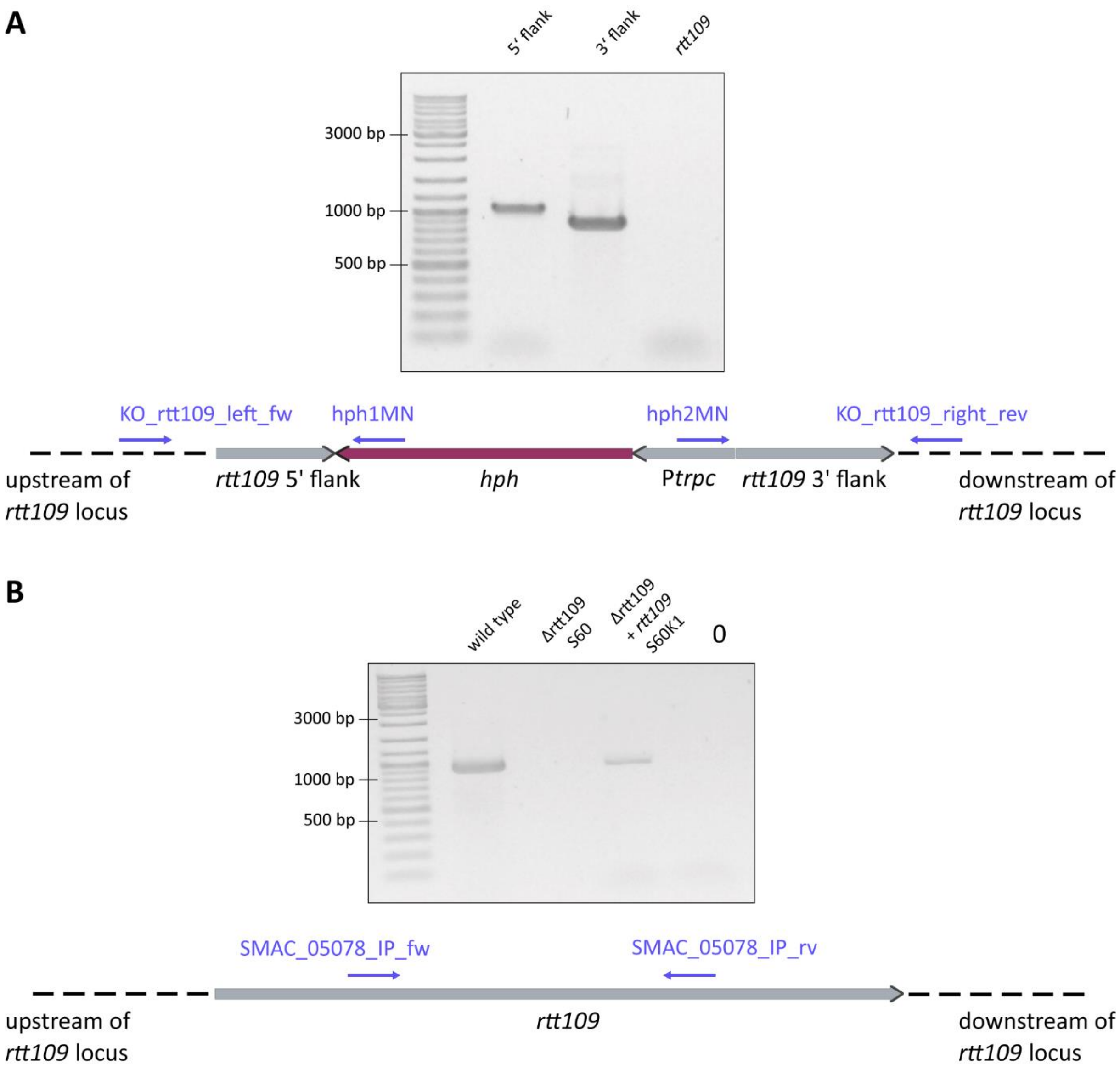


**Figure S1:** Confirmation of *rtt109* deletion and complementation by PCR. Examples of agarose gels for Δrtt109 strain S60 and complementation strain S60K1 are shown. PCR primers are given as blue arrows.
**A.** Verification of the correct integration of the *rtt109* deletion cassette in *S. macrospora* Δrtt109 S60. Amplification of a part of the *rtt109* upstream region and the integrated *hph* sequence should yield a PCR product of 1050 bp, while the downstream region should yield a product of 920 bp. Bands of the expected size were detected by agarose gel electrophoresis.
**B.** Amplification of an internal part of the *rtt109* gene should produce a PCR product of 932 bp, which was not detected in the mutant. Successful reintroduction of *rtt109* into *S. macrospora* Δrtt109 should yield PCR products of 932 bp, indicating the presence of *rtt109*. The expected band was detected in the wild type and the complementation strain, but not in the deletion mutant.

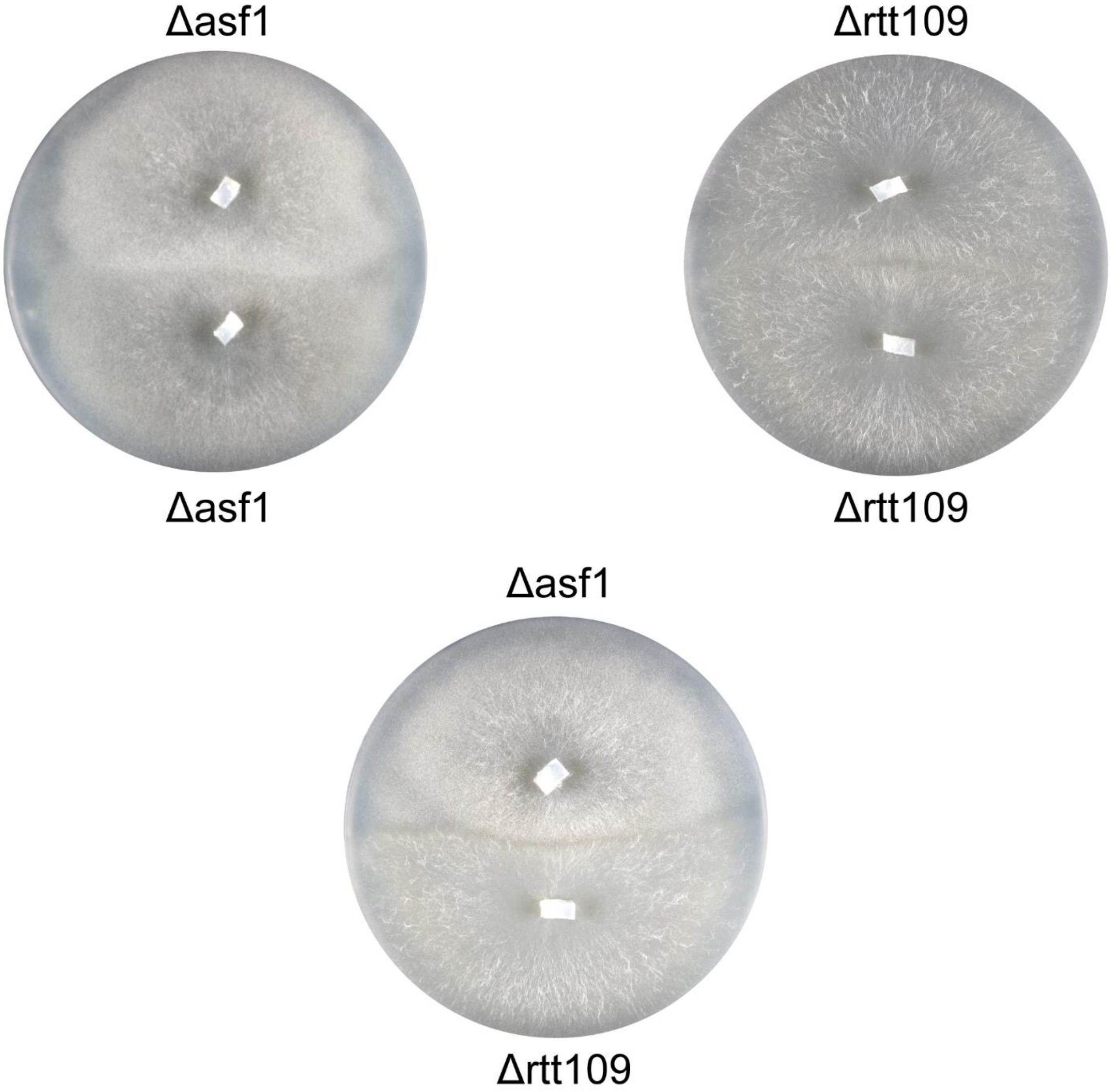


**Figure S2.** Crossing attempt between *S. macrospora* Δasf1 and Δrtt109 strains. The mutants do not produce fruiting bodies in the contact zone. As a control, the Δasf1 and Δrtt109 strains were also crossed against themselves, and these crosses are sterile and do not form fruiting bodies (as expected). The strains were co-cultured on BMM medium for 8 days.

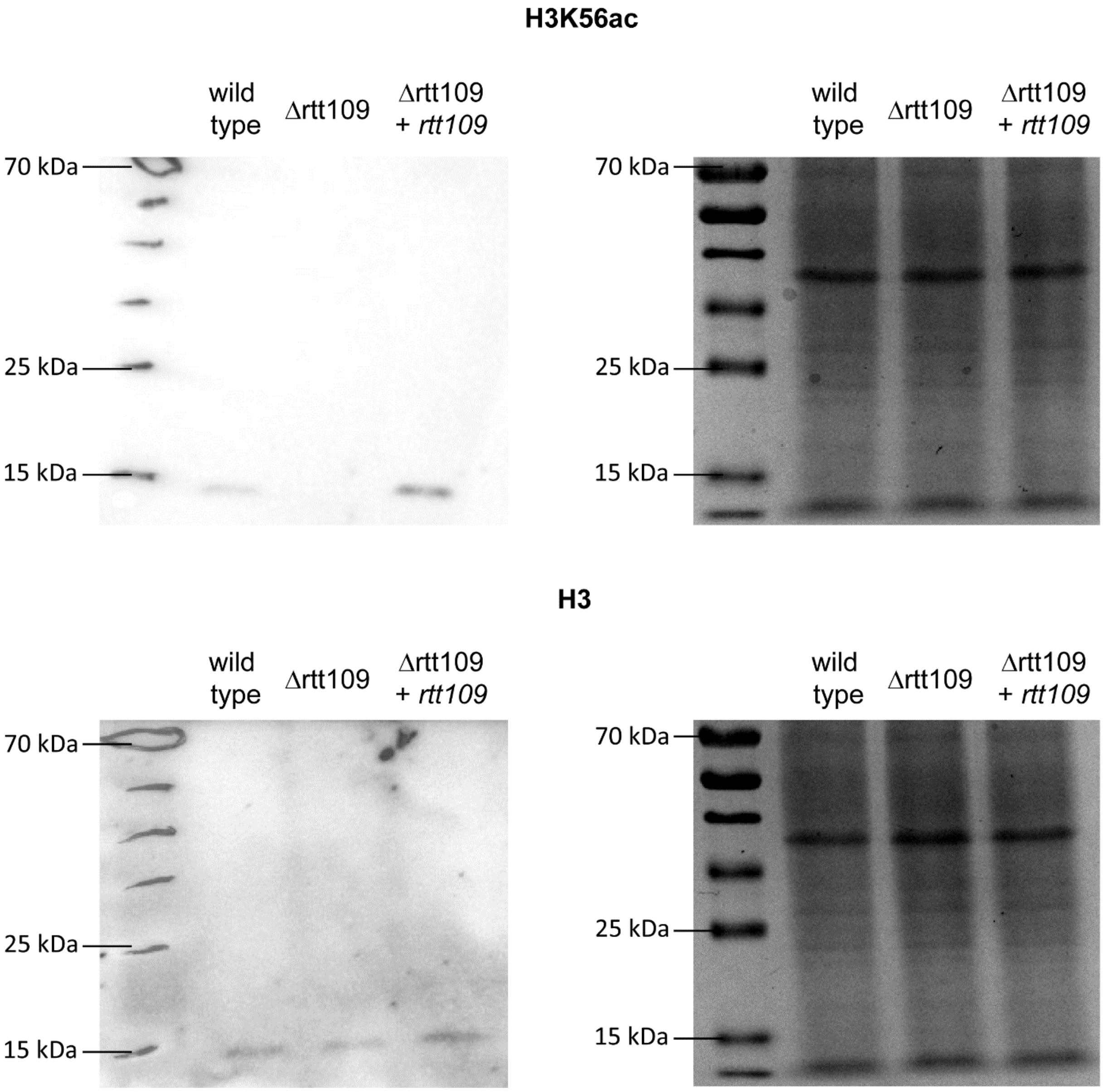


**Figure S3.** Uncropped Western blot (left) and Coomassie-stained gel (right) for assessment of H3K56ac and H3 levels. Protein extracts from the wild type, rtt109 deletion mutant and the respective complementation strain were used as samples in western blot experiments. H3K56ac and H3 signals were detected by antibodies specific for the modification H3K56ac (top) and histone H3 (bottom).

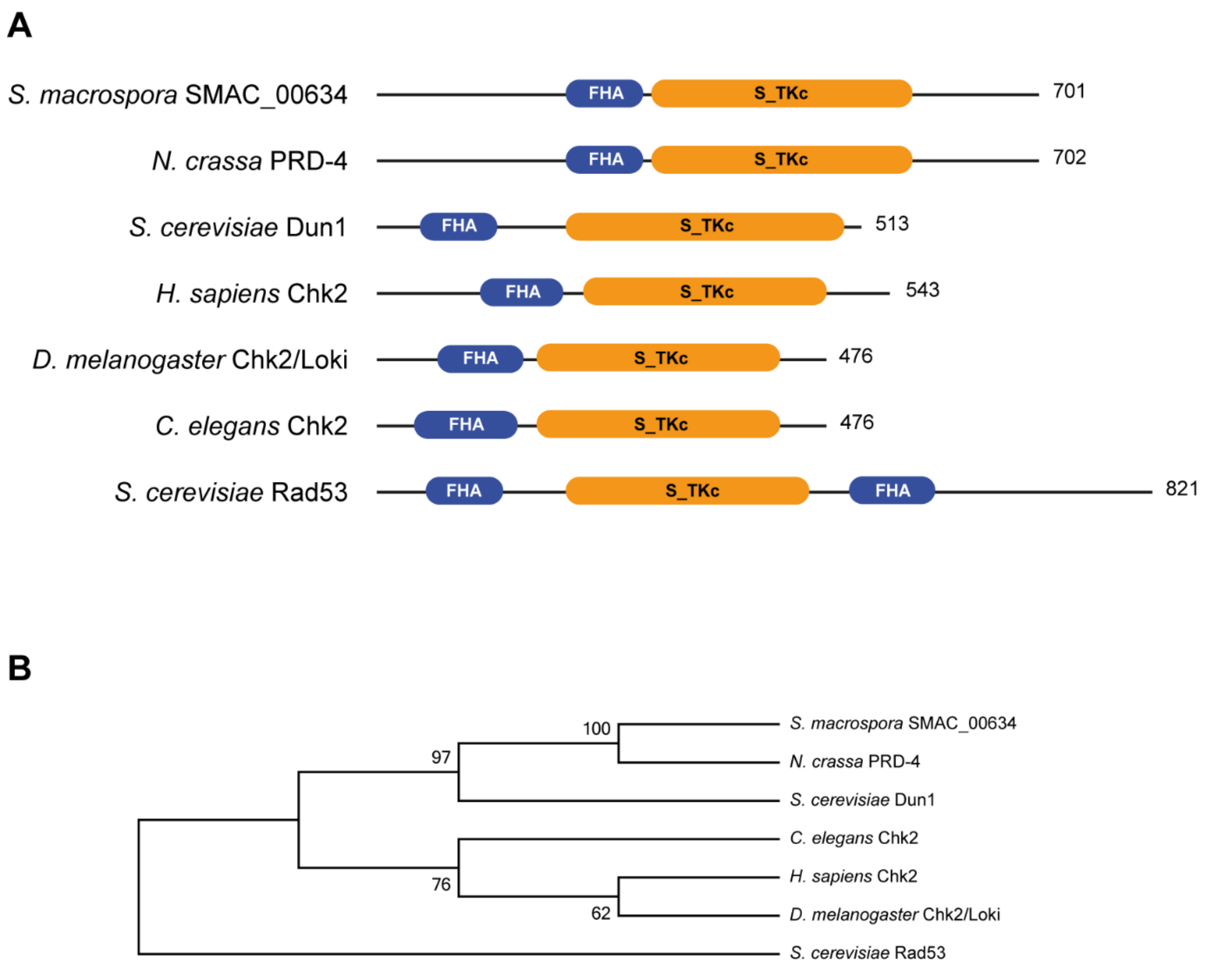


**Figure S4.** RAD53 homologs in fungi and animals.

**A.** Domain structure of RAD53 homologs. The length of the proteins in amino acids is given on the right. FHA: forkhead-associated domain, S_TKc: catalytic domain of serine/threonine protein kinase.

**B.** Phylogenetic tree of RAD53 homologs. Proteins sequences were aligned with ClustalX and a phylogenetic tree was calculated in Mega6 with Neighbor joining (1000 bootstrap replicates, bootstrap percentages are indicated at the branches). The following sequences were used for the analysis: *Sordaria macrospora* SMAC4_00634 (CHK2), *Neurospora crassa* PRD-4 XP_964470.3, *Saccharomyces cerevisiae* Dun1 YDL101C SGDID:S000002259, *Homo sapiens* Chk2 NP_009125.1, *Drosophila melanogaster* Chk2/Loki NP_477219.1, *Caenorhabditis elegans* Chk2 NP_001024271.1, *Saccharomyces cerevisiae* Rad53 YPL153C SGDID:S000006074.

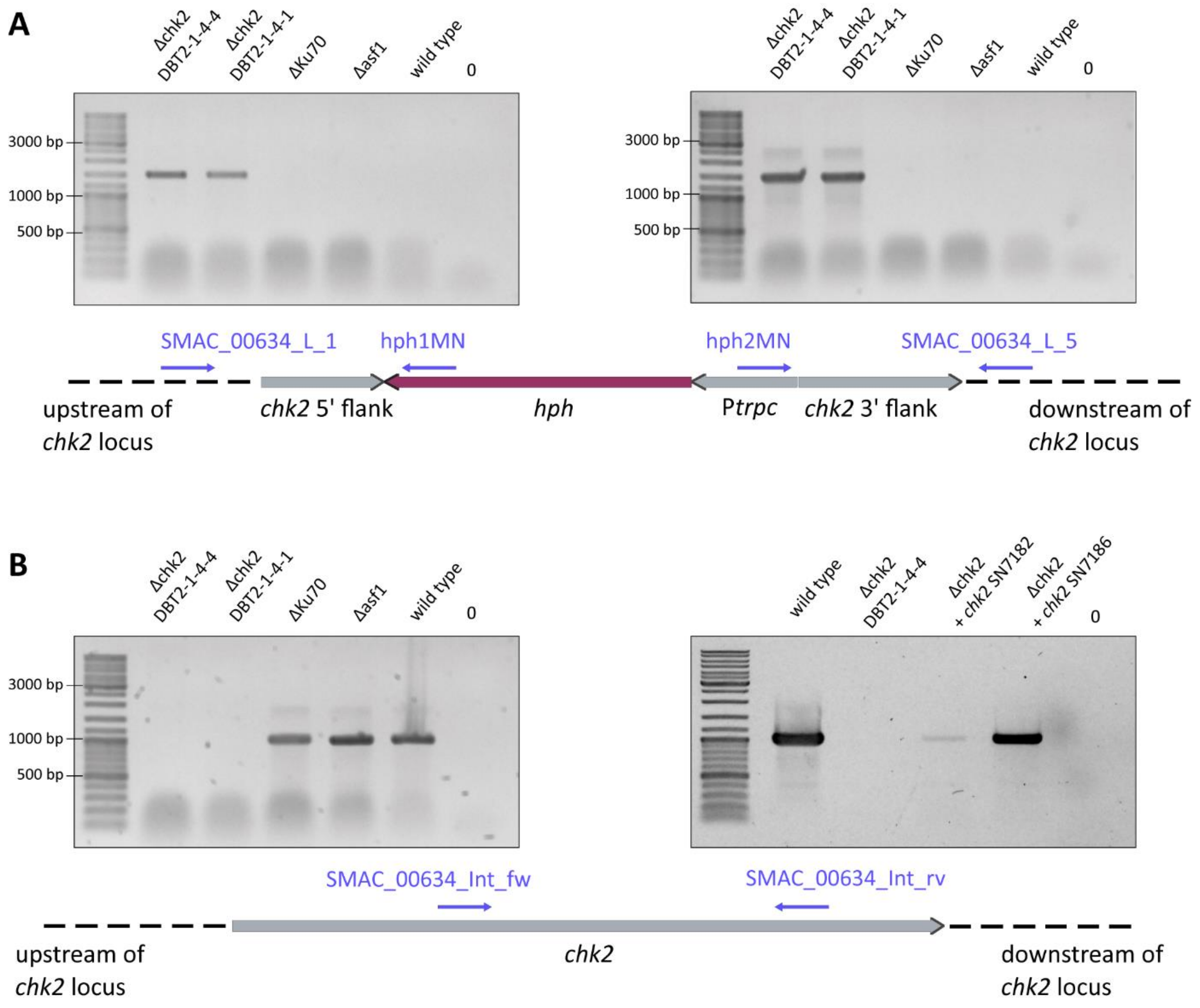


**Figure S5.** Confirmation of *chk2* deletion and complementation by PCR. Examples of agarose gels for Δchk2 strains DBT2-1-4-4 and DBT2-1-4-1 are shown. PCR primers are given as blue arrows.

**A.** The expected PCR product of 1460 bp for amplification of the *chk2* upstream region and integrated *hph* sequence and 1529 bp for the downstream region, indicating correct integration of the *chk2* deletion cassette, was detected in the tested mutants.

**B.** PCR with primers designed for the amplification of an internal part of *chk2* yielded the expected product of 1028 bp in all strains, except the deletion mutants.

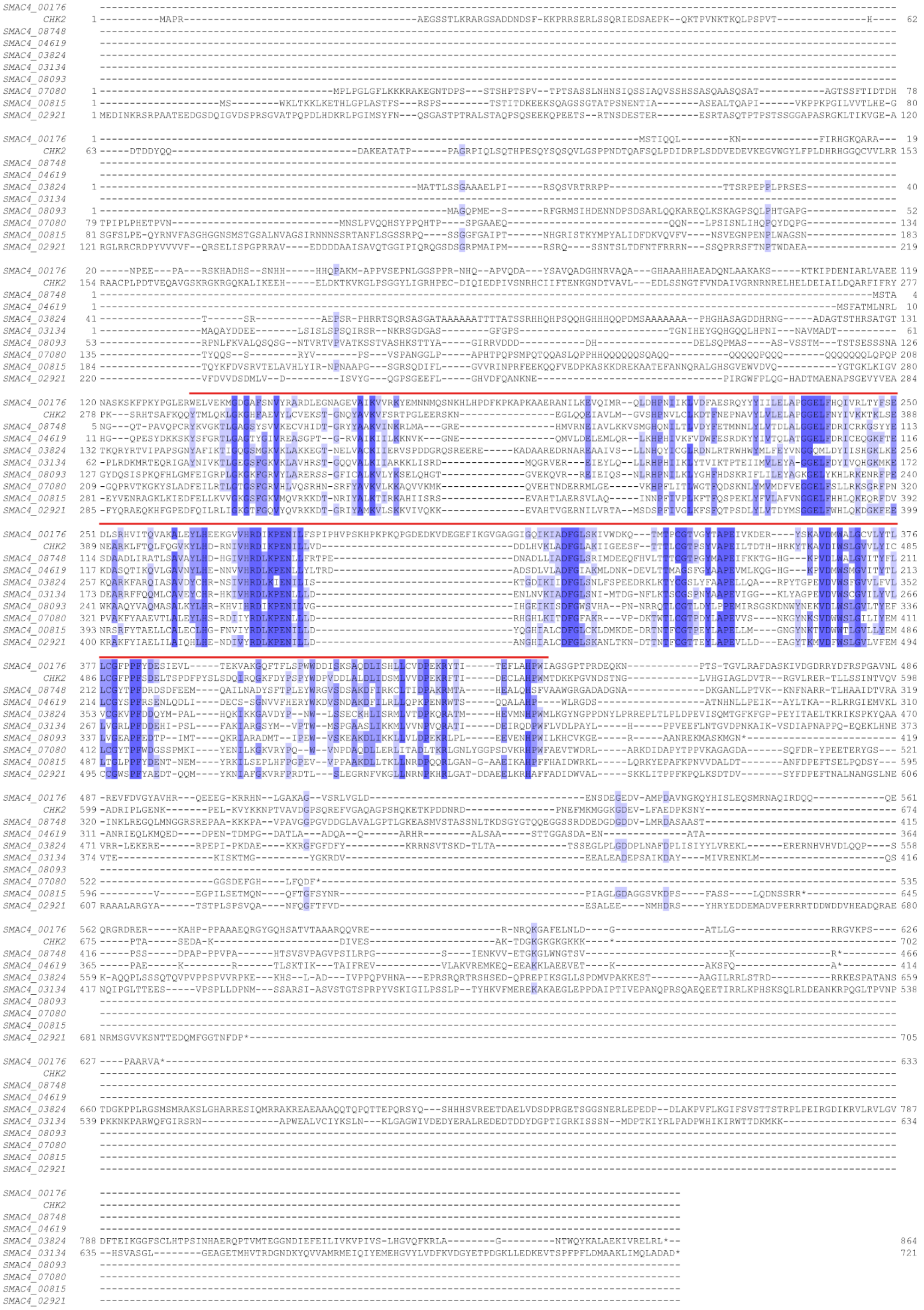

**Figure S6.** Multiple alignment of BLASTP hits (evalue < 0.1) of CHK2 among the predicted *S. macrospora* proteins. The S_TKc domain (Serine/threonine protein kinase, catalytic domain) present in all proteins is indicated with a red line above the sequences. Apart from the S_TKc domain, the BLASTP hits do not have regions of high similarity to CHK2. The alignment was generated with Clustal Omega [Madeira et al. (2022), Nucleic Acids Research, 50(W1):W276-W279].

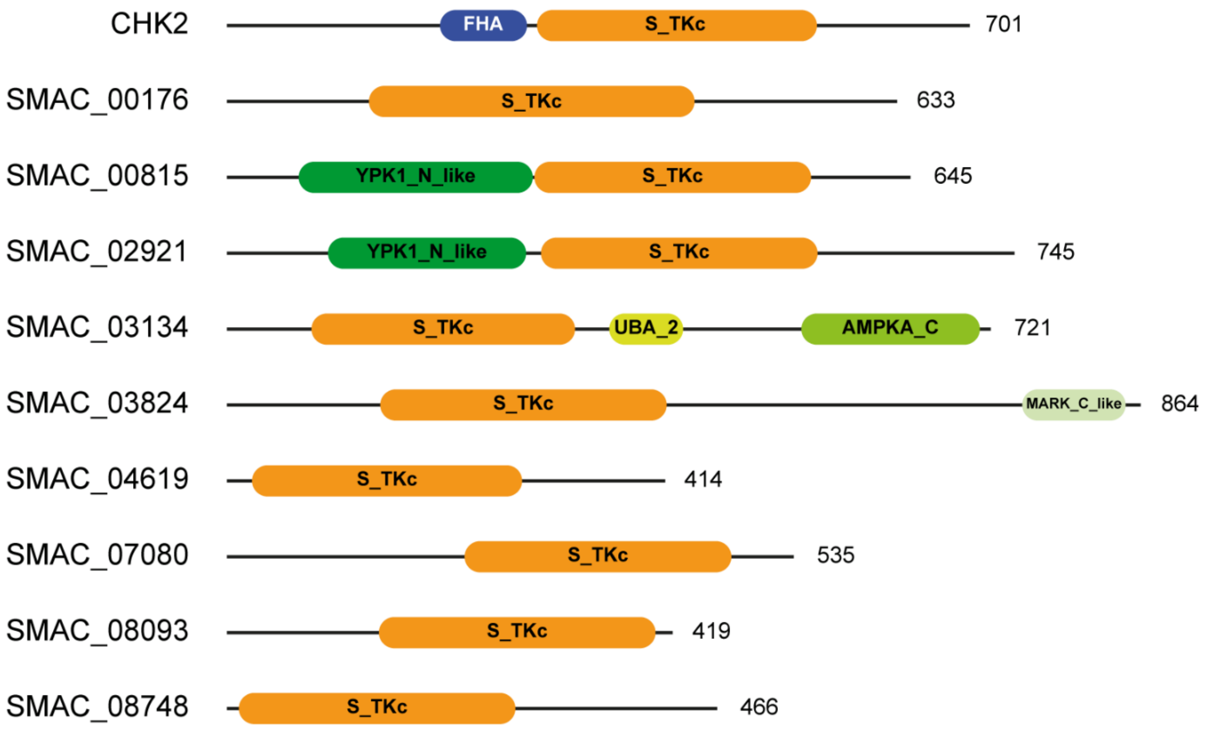


**Figure S7.** Domain structure of BLASTP hits (e-value < 0.1) from a search with CHK2 (SMAC_00634) among the predicted *S. macrospora* proteins. The length of the proteins in amino acids is given on the right. None of the BLASTP hits has the same domain structure as CHK2, i.e. an S_TKc domain combined with an FHA domain. Domain abbreviations: AMPKA_C: C-terminal regulatory domain of 5'-AMP-activated protein kinase alpha catalytic domain, FHA: forkhead-associated domain, MARK_C_like: C-terminal kinase associated domain 1, S_TKc: catalytic domain of serine/threonine protein kinase, UBA_2: Ubiquitin-associated domain, YPK1_N_like: Fungal protein kinase domain similar to the N-terminus of YPK1.
